# A continuum of asynchronous states in cerebral cortex networks, and how they determine responsiveness

**DOI:** 10.64898/2026.05.06.723408

**Authors:** Matthew Bassat, Federico Tesler, Alain Destexhe

## Abstract

The awake brain is known to display asynchronous (AS) states during periods of attention and arousal, but the responsiveness properties of such states remain unclear. Here, we investigate this question using computational models of spiking networks of excitatory and inhibitory neurons, mimicking recurrently-connected networks in layer 2/3 of the cerebral cortex. The networks can generate a continuum of AS states, but with different responsiveness characteristics. By using a mean-field model to infer the dynamic properties of the system, we find that there are two families of AS states, which we call “underdamped” (UD) and “overdamped” (OD). Responsiveness is maximised at the transition between OD and UD states, which identifies a “working point” that may present advantageous computational properties.

## 1 Introduction

The awake cortex is typically associated with asynchronous (AS) firing activity, characterised by stable population firing rates and low neuronal synchrony [1–3]. At the whole brain scale, cortical responses in AS states have been shown to amplify external inputs relative to other brain states, chiefly the Up and Down regime associated with slow-wave sleep (SWS) and anaesthesia [4]. A pioneering work by Massimini et al. [5] showed that cortical electroencephalographic (EEG) responses to transcranial magnetic stimulation (TMS) in humans are longer-lasting, more diffuse and more complex in the awake state than in unconscious brain states. At the single neuron and local network scales, the high-conductance state typical of AS networks has been shown to confer enhanced computational capacity and enhanced responsiveness to low-amplitude inputs [6]. A natural corollary is to study the correlates of responsiveness within the AS regime, and to ask under what conditions AS states are most responsive.

Following Ledoux and Brunel [7], who thoroughly characterised the responsiveness of cortical AS networks to time-dependent afferent impulses in analytically tractable models (networks of fully-connected leaky integrate-and-fire neurons with current-based synapses, and Wilson–Cowan rate models with threshold-linear transfer functions), we address this question in a more biologically realistic spiking network (SNN) model. Namely, in sparsely-connected networks of adaptive exponential integrate-and-fire (AdEx) neurons [8] with conductance-based (COBA) synapses. From these elements, we model layer 2/3 (L2/3) cortical circuits comprising excitatory regular-spiking (RS) and inhibitory fast-spiking (FS) populations (see 4) and interrogate the responsiveness of these networks under different parameter conditions. Despite its simplicity, consisting of only two differential equations, the AdEx model is known to reproduce the activities of cortical RS and FS cells to high accuracy [8] with biologically interpretable parameters. Further, the COBA synaptic model ensures that the conductance state of the network is taken into account in the integration of synaptic inputs [9, 10], which is of particular relevance in a study of responsiveness. To emphasise the generality of our findings, we show that the qualitative results obtained from AdEx simulations are reproduced in the more biologically realistic Hodgkin-Huxley (HH) model [11].

Several studies have identified different classes of AS states with distinct responsiveness characteristics. In this paper, we take a dynamical systems approach, treating AS states as two-dimensional stable fixed points and classifying them based on their linear stability; fixed points with negative real eigenvalues and complex conjugate eigenvalues are termed overdamped (OD) and underdamped (UD), respectively. Previously, AS networks have been classified into *recurrent input-dominated* and *afferent-dominated* categories based on the level of background excitatory drive [12], conferring relatively high responsiveness and strong pattern-encoding properties, respectively. Another study defined AS states as *homogeneous* or *heterogeneous* based on the level of synaptic coupling between and within populations [13], with the former favouring signal transmission (through redundant, low-dimensional responses) and the latter providing a richer substrate for complex computations (through higher-dimensional responses). A third study measured the responsiveness of AS states as a function of the level of heterogeneity in the parameters, finding that responsiveness was optimised at experimentally observed levels of heterogeneity [14]. While these approaches identify meaningful functional categories, we argue that our categorisation provides a complementary and practically useful framework, as it centres on a universal transition between regimes that emerges as the E/I balance in the network is modulated. In addition, our approach reveals a maximum in responsiveness to slow inputs at a “working point” occurring at the boundary of the two regimes, making this a doubly useful categorisation method.

By investigating the spiking network and its mean-field in parallel, we show that the mean-field reduction provides a simple and effective analytical framework for predicting the responsiveness of AS states.

## 2 Results

Responsiveness simulations were conducted on artificial L2/3 cortical circuits (Fig. 1a) modelled by sparsely-connected spiking neural networks of excitatory RS and inhibitory FS neurons, using the AdEx model (see 4.1 for details). The E/I balance of the network was tuned via changes in 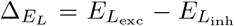 (the difference between excitatory and inhibitory leak potentials), and a range of parameter combinations yielding AS activity (stable population firing rates) was identified.

**Fig. 1:**
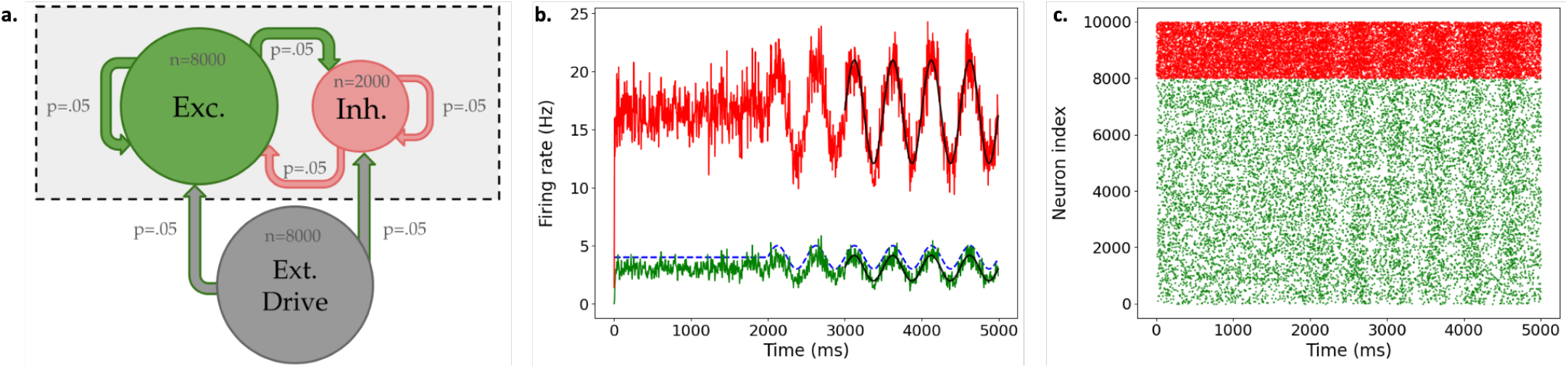
**(a)** Schema of L2/3 spiking network model, consisting of 8000 excitatory neurons, 2000 inhibitory neurons, and an external population of 8000 excitatory Poisson neurons supplying a background drive to both populations. All neurons are randomly connected, with uniform connection probability (0.05). **(b)** Population firing rate plot from example responsiveness simulation (*ν*_drive_ = 4 Hz, *b* = 0 pA, 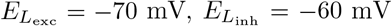, input amplitude *A* = 1 Hz, input frequency *f* = 2 Hz). Green (red) traces: mean excitatory (inhibitory) firing rate (5 ms bins). Blue dashed line: time-varying firing rate of external input. Black lines: sinusoidal fits of evoked population responses. **(c)** Raster plot from responsiveness simulation in **b** (10% of spikes shown for visibility).

The linear stability of an asymptotically stable fixed point in a two-dimensional dynamical system (i.e. the eigenvalues of its jacobian matrix) can fall into one of two categories: a fixed point with two negative real eigenvalues (either distinct or repeated) is a *stable node*, while a fixed point with complex conjugate eigenvalues (with negative real parts) is a *stable focus*. Here, we borrow the terms overdamped (OD) and underdamped (UD) from the physics of damped harmonic oscillators to describe fixed points with negative real eigenvalues and complex conjugate eigenvalues, respectively. A mean-field model [15] derived from the spiking network (see 4.3) was used to obtain a low-dimensional description of the network activity, represented by two dimensions (mean excitatory and inhibitory population firing rates), and to numerically analyse its linear stability as a two-dimensional system. The mean-field model has an associated timescale *T*, which significantly affects eigenvalue calculations; in a departure from previous studies, a state-dependent timescale was used to fix a theoretically minimal, non-arbitrary choice of *T* for a given parameter combination (see 4.3.4 for details). For the range of AS states explored in the spiking network, the eigenvalues of the associated mean-field networks were computed, following the method outlined in 4.3 (Fig. 2a).

**Fig. 2:**
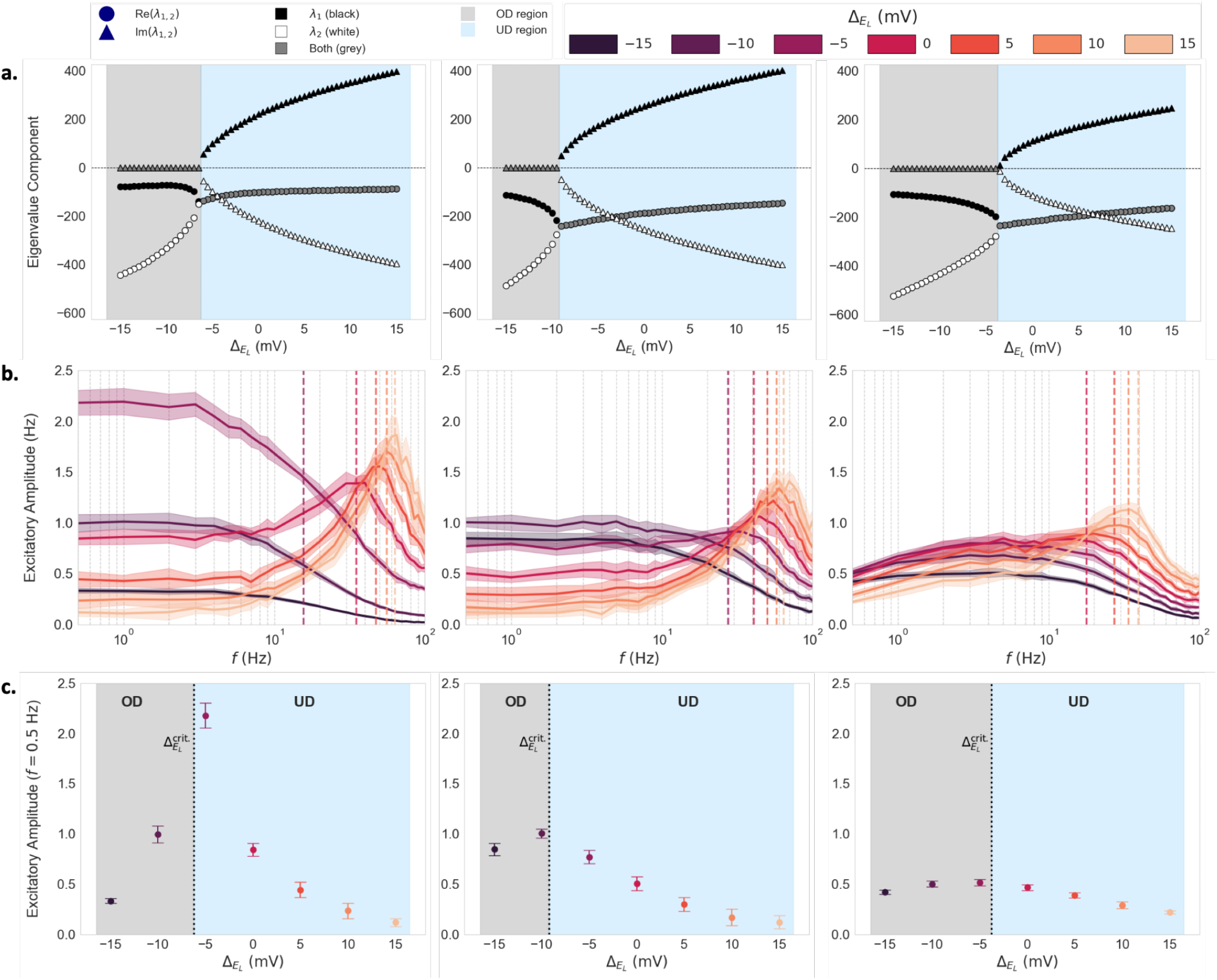
Mean-field linear stability analysis and SNN frequency-response profiles for different 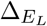 values 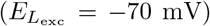 across parameter combinations (input amplitude *A* = 1 Hz). **First column**: *b* = 0 pA, *ν*_drive_ = 2 Hz; **second column**: *b* = 0 pA, *ν*_drive_ = 4 Hz; **third column**: *b* = 60 pA, *ν*_drive_ = 4 Hz. Eigenvalues versus 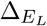. Shapes indicate real (circles) and imaginary (triangles) parts. Colours denote eigenvalue index (black: *λ*_1_; white: *λ*_2_; dark grey: common to both). Background colours denote OD (light grey) and UD (blue) regimes. **(b)** Amplitudes of evoked excitatory responses versus input frequency *f* ∈ [0.5, 100] Hz (log scale), coloured by 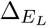. Dashed lines: eigenfrequencies for UD subset. **(c)** Visualisation of evoked responses to a slow input (*f* = 0.5 Hz). 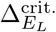 denotes the mean-field prediction of the boundary between OD (grey) and UD (blue) regions.

To measure their responsiveness, networks were subjected to time-dependent sinusoidal impulses following Ledoux and Brunel [7] and Di Volo et al. [15] (see example in Fig. 1b and c), with evoked responses quantified by the amplitudes of the evoked changes in the average population firing rates, per 4.2 (we later explore the responses to non-oscillating inputs in 2.4). Responses were measured across a broad range of input frequencies (0.5 ≤ *f* ≤ 100 Hz, amplitude fixed at *A* = 1 Hz), with the results presented in Fig. 2b (equivalent results for *A* = 2 Hz are shown in Supplementary Fig. S1). In Fig. 2c, the cross-section of responses to the slowest input (*f* = 0.5 Hz) is shown as a function of 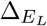.

### 2.1 Overdamped and underdamped regimes: low-pass and resonant response profiles

Across the range of parameters explored, the frequency-response profiles (Fig. 2b) were robustly divided into two distinct categories along the 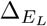 axis. For low 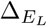 values, the response curves show low-pass behaviour (disregarding adaptation effects for low frequencies, see 2.3). When 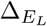 exceeds some threshold, resonances appear in the beta and low gamma ranges (∼12-70 Hz).

#### 2.1.1 Capture of SNN responsiveness characteristics by linear stability analysis

To understand the emergence of the two different regimes, we make use of the meanfield model of the cortical circuit. We show here that the dichotomy of responsiveness profiles in the SNN model is comprehensively explained by the mean-field linear stability analysis, with low-pass and resonant networks corresponding to stable nodes (OD states) and stable foci (UD states), respectively.

For the range of AS networks studied, the eigenvalues of the jacobian matrices of the corresponding mean-field representations are shown in Fig. 2a. The 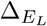 values at which the response curves transition from low-pass to resonant (Fig. 2b) are well predicted by the nature of the computed eigenvalues, with the former behaviour corresponding to real-eigenvalued fixed points and the latter corresponding to complex-eigenvalued fixed points, as observed in Ledoux and Brunel [7]. With the exception of 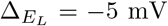 in the first parameter region (Fig. 2b first column), which is misidentified as a resonant network (due to its proximity to the estimated damping boundary), the responsiveness categories of all simulated networks are correctly classified by the eigenvalue analysis.

Secondly, the resonant frequencies in UD networks are well captured by the meanfield analysis. Such networks are close to a supercritical Hopf bifurcation (at Re(*λ*_1,2_) = 0), where population firing rates begin to oscillate periodically due to the interplay between excitatory and inhibitory synaptic time scales [16]. Their resonant frequencies are theoretically predicted by the eigenfrequencies of the associated stable foci [7]:

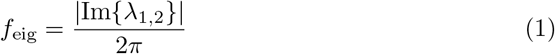

In Fig. 2b, the dashed lines indicate the eigenfrequencies corresponding to the plotted 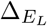 values for the states classified as UD. We observe that, across the three distal parameter regions, and spanning a broad frequency range, the positions of the responsiveness peaks are well predicted by the mean-field eigenfrequencies, albeit with reduced precision for high *b* (see 2.3).

Thirdly, the amplitudes of the UD resonance peaks are strongly inversely correlated with |Re {*λ*_1,2_ }, the moduli of the real parts of the complex conjugate eigenvalues (Fig. 3). In other words, the peak heights are proportional to the proximity of the network to the Hopf bifurcation line in the mean-field prediction (as in Ledoux and Brunel [7]), and are therefore inversely proportional to the stability of the network.

**Fig. 3:**
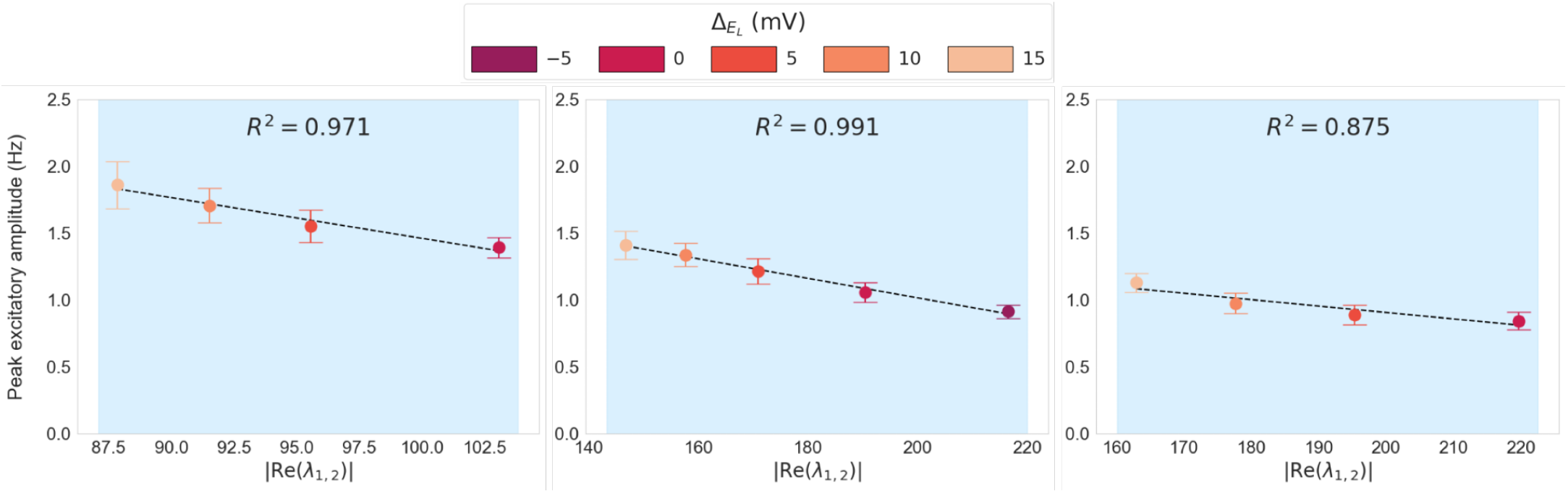
Maximum evoked excitatory response amplitude (resonance peak height) versus modulus of real part of complex conjugate eigenvalues, |Re(*λ*_1,2_)|, across 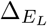 values in the UD region. Dotted lines: line of best fit. **Columns 1-3**: parameter sets as in Fig. 2. Note the exclusion of 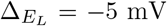 from column 1 (misidentified as a UD network).

#### 2.1.2 Overdamped regime: responsiveness is correlated with excitation-dominance for all frequencies

In all parameters regions explored, the evoked response amplitudes correlate positively with 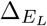 within the OD regime across all input frequencies. Networks in this regime exhibit no resonances and evoked responses monotonically increase as input frequency decreases until saturating below a certain threshold (when the input is significantly slower than the relaxation time of the network, and firing rates are at equilibrium as the input varies, i.e., a quasi-stationary regime). In the low-frequency limit, the response depends on the competition of excitatory and inhibitory gains and increases as 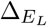 approaches the working point (i.e. the damping boundary) from the left (as discussed in 2.2). Therefore, cortical networks in the OD regime are in a generalised, indiscriminate responsiveness mode in the low frequency range, with the degree of responsiveness determined by the proximity of the network to the working point (see Fig. 2c, OD regions).

#### 2.1.3 Underdamped regime: away from the resonance peak

In the low-frequency limit, the amplitude of the responses in the UD regime decrease as 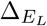 increases, opposite to the behaviour in the OD regime. For sufficiently fast inputs (*f* ≫ *f*_eig_), responses are consistently proportional to 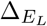 (Fig. 2b) across both damping regimes, scaling with the stationary population firing rates (which scale with 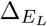, see Supplementary Information SI.2), consistent with the high-frequency behaviour of the transfer function in networks of leaky integrate-and-fire neurons [7].

### 2.2 Responsiveness to slow inputs peaks at the damping regime boundary, 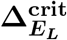

Across parameter sets, slow-input responsiveness for both excitatory and inhibitory populations is maximised near the transition 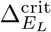 between overdamped and underdamped modes (Fig. 2c).

In this section, we derive an analytical estimate of the responsiveness of the network from the mean-field equations. The mean-field model used here is built upon a semi-analytic transfer function *F*_*µ*_ [15, 17], which describes the output firing rate of a neuron in population *µ* ∈ {*e, i*} as a function of its excitatory and inhibitory presynaptic input rates, *ν*_*e*_ and *ν*_*i*_ (see 4.3.2 for details).

For a given AS fixed point 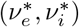 with a baseline excitatory drive of *ν*_drive_ Hz, the excitatory and inhibitory firing rates are approximated by the transfer function estimates 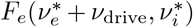 and 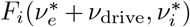, respectively (fixing the excitatory adaptation current *w* at its stationary value).

We estimate the magnitudes of the evoked firing rate deviations *δν*_*e*_ and *δν*_*i*_ to a small increase *δν*_drive_ in the excitatory drive by evaluating the excitatory (inhibitory) transfer function at 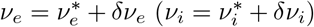 with a first-order Taylor expansion, obtaining the following expression (see Supplementary Information SI.3 for derivation):

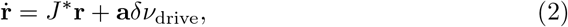

where **r** = (*δν*_*e*_, *δν*_*i*_)^⊤^, *J* ^∗^ ∈ ℝ^2×2^ is the jacobian matrix (4.3.3) and **a** is the vector 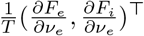 evaluated at the fixed point in question.

For sufficiently slow inputs, that is, much slower than the relaxation time constant of the network, which is on the order of *T* = *τ*_*V*_ ≈ 10 ms (see 4.3.4), we may make the quasi-stationary assumption that the time derivative of the changes in firing rates 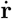 is approximately zero, that is, that the firing rates are stationary for a given input. This gives the following linear relationship between the afferent perturbation and the evoked responses:

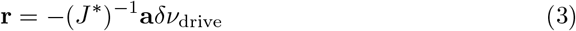

Expressed in terms of the notation in 4.3.3, we obtain the following expression:

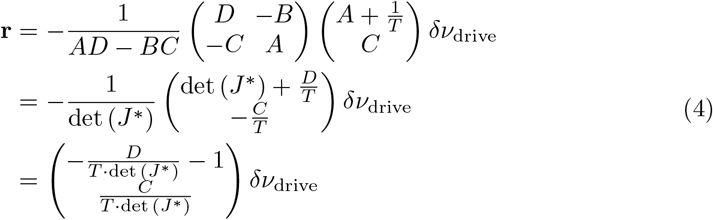

A comparison of the evoked excitatory and inhibitory responses to a slow input (*A* = 1 Hz, *f* = 0.5 Hz) from SNN simulations against the analytical estimates *δν*_*e*_ and *δν*_*i*_ (fixing *δν*_drive_ = 1 Hz) is shown in Figure 4. We observe that the linearisation predicts a point of maximal responsiveness to slow inputs for both excitatory and inhibitory populations near the working point, consistent with simulations.

**Fig. 4:**
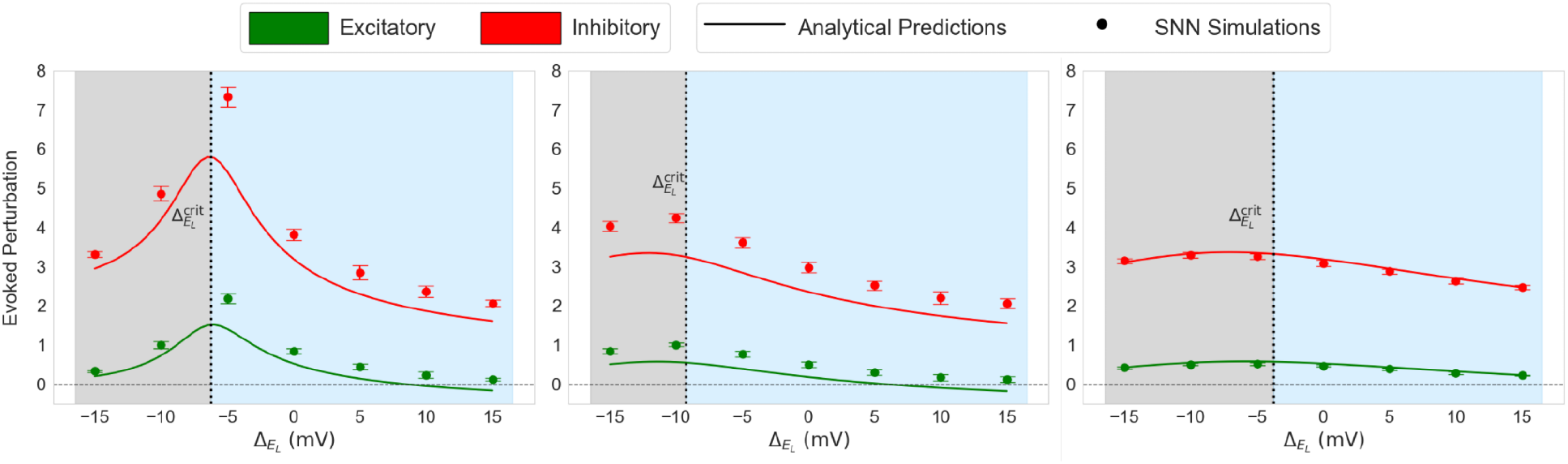
Analytical estimates *δν*_*e*_ and *δν*_*i*_ (solid lines) for the excitatory (green) and inhibitory (red) evoked perturbations induced by an arbitrarily slow increase in afferent drive *δν*_drive_ = 1 Hz as a function of 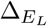, overlaid with SNN network responses (circles) to a slow oscillatory input (*A* = 1 Hz, *f* = 0.5 Hz). Grey (blue) represents the overdamped (underdamped) region. **Columns 1-3**: parameter sets as in Fig. 2.

### 2.3 Changing adaptation and external drive

Increasing the excitatory spike adaptation current *b* from 0 to 60 pA (Fig. 2, columns 2 and 3) shifted 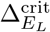 to the right, substantially lowered the resonant frequencies of UD networks and slightly lowered the resonance peak amplitudes, consistent with the substantial associated decreases in |Im{*λ*_1,2_}| (Fig. 2a) and subtle increases in |Re {*λ*_1,2_} (Fig. 3). Therefore, depending on the excitability of the system, an increase in spike-dependent adaptation can either induce a transition from a resonant to a low-pass responsiveness regime, or can strongly modulate the preferred frequency of a network within the resonant regime (see Supplementary Information SI.4 for a more extensive exploration of changes in *b*). The level of spike-frequency adaptation in pyramidal neurons is subject to neuromodulation, e.g. by acetylcholine [18], providing a possible mechanism for adaptive frequency selectivity.

For non-zero *b*, responses began to decrease as input frequency decreased at very low frequencies due to the slow accumulation of adaptation (Fig. 2b third column), which evolves at a very slow timescale (*τ*_*w*_ = 500 ms). This effect only emerges for inputs on the order of the adaptation timescale [15], i.e. 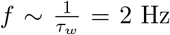. Moreover, the size of this effect is approximately proportional to 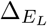, since, for a given *b*, the mean accumulation in adaptation *w* is proportional to the mean firing rate, which scales with 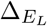.

Increasing *ν*_drive_ from 2 to 4 Hz (Fig. 2, first and second columns) shifted 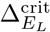 to the left and substantially diminished the heights of the resonance peaks in UD networks, as well as minimally increasing the resonant frequencies. This is well captured by the uniform increase in |Re{*λ*_1,2_}| and relatively unchanged |Im{*λ*_1,2_}| in the UD region as *ν*_drive_ increases (Fig. 2a, columns 1 and 2). The same effects were observed when the drive was increased to 8 Hz (see Supplementary Information SI.5). As a consequence, resonant networks can maintain their intrinsic frequency selectivities in the presence of substantial changes in the level of background input from neighbouring cortical regions, albeit with reduced peak responses for increasing drive.

More broadly, increasing *ν*_drive_ flattened the frequency-response curves across both regimes: the peak response at the working point was reduced and the slow-input response curve became shallower, a consequence of the relatively large determinant at 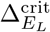, and the relatively gentle rates of change in the excitatory and inhibitory gains, for higher background drive (see Supplementary Information SI.6). Since OD responses decay monotonically from their low-frequency maximum, this flattening extends across all frequencies for OD states. Therefore, while increasing background drive can shift a network from the OD to the UD regime, its primary effect in both regimes is a general reduction in responsiveness across input frequencies. One exception to this is the responsiveness of OD networks near the damping boundary, which in some cases can increase for larger external drive (e.g. 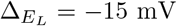, see Fig. 2c) as a consequence of the flattened response curves, despite the lower peak response.

### 2.4 Responses to pulse inputs

To verify whether our results generalised to pulse-type (non-oscillating) inputs, we ran alternative responsiveness simulations with sinusoidal half-cycle inputs [15] at a range of timescales for representative OD, working point and UD networks.

As in the case of oscillatory inputs, the network near the working point (Fig. 5b) showed markedly higher responses than both the OD and UD networks to the slow (first column) and moderate (second column) impulses. The UD network showed weak responses to both of these inputs, especially in the excitatory population; interestingly, it did not show a significantly larger response to the fast input (third column), with the response barely exceeding the baseline noise, despite the fact that the input timescale was equivalent to the resonant frequency of the network (Fig. 2b, first column). Therefore, while the working point result is replicated for single pulse inputs, the high-frequency resonances for UD states effectively disappear.

**Fig. 5:**
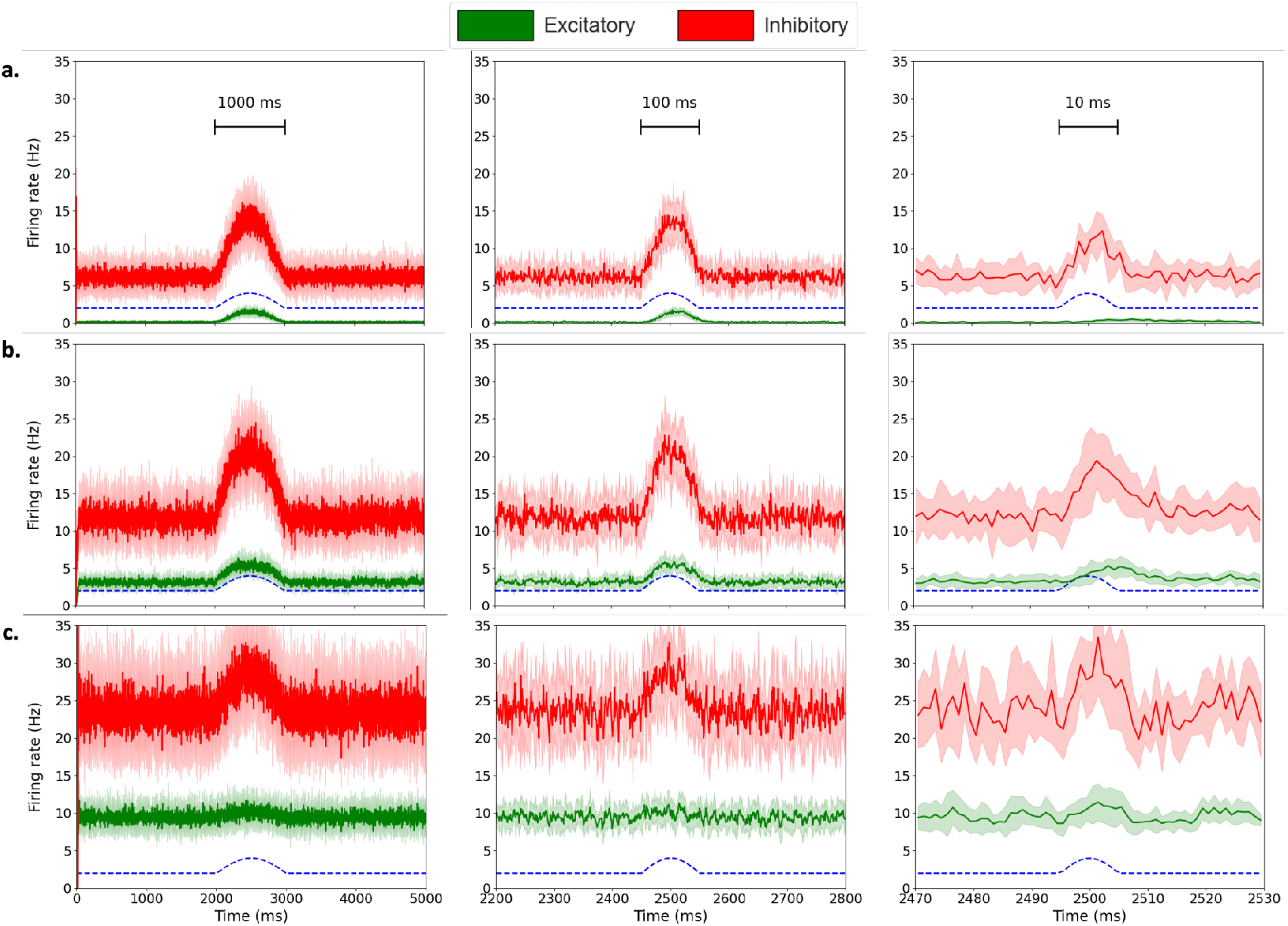
Sinusoidal half-cycle responsiveness simulations (averaged across 10 seeds, 1ms bins, shading indicates std across seeds) across overdamped, working point and underdamped networks for slow, moderate and fast input frequencies (input amplitude fixed at *A* = 2 Hz). **(a)** OD network (*b* = 0 pA, *ν*_drive_ = 2 Hz,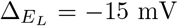). **(b)** Network close to working point (*b* = 0 pA, *ν*_drive_ = 2 Hz,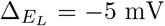). **(c)** UD network (*b* = 0 pA, *ν*_drive_ = 2 Hz,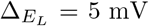). **Column 1:** Half-cycle length: 1000 ms (equivalent frequency: *f* = 0.5 Hz). **Column 2:** Half-cycle length: 100 ms (equivalent frequency: *f* = 5 Hz). **Column 3:** Half-cycle length: 10 ms (equivalent frequency: *f* = 50 Hz, close to the resonant frequency of network **c**).

In Fig. 6, we repeat the UD simulations but replace the pulse input with two full oscillation periods. The large augmentation of the response to the fast input (third column) only emerges after the first half-cycle, requiring additional time for the network to reach its maximal amplitude. It follows from this observation that the resonant properties of UD states only occur in response to a sustained oscillation (of at least one full period). This reaffirms the dual functional roles of OD and UD states, with the latter being specifically responsive to oscillations at or near their resonant frequencies.

**Fig. 6:**
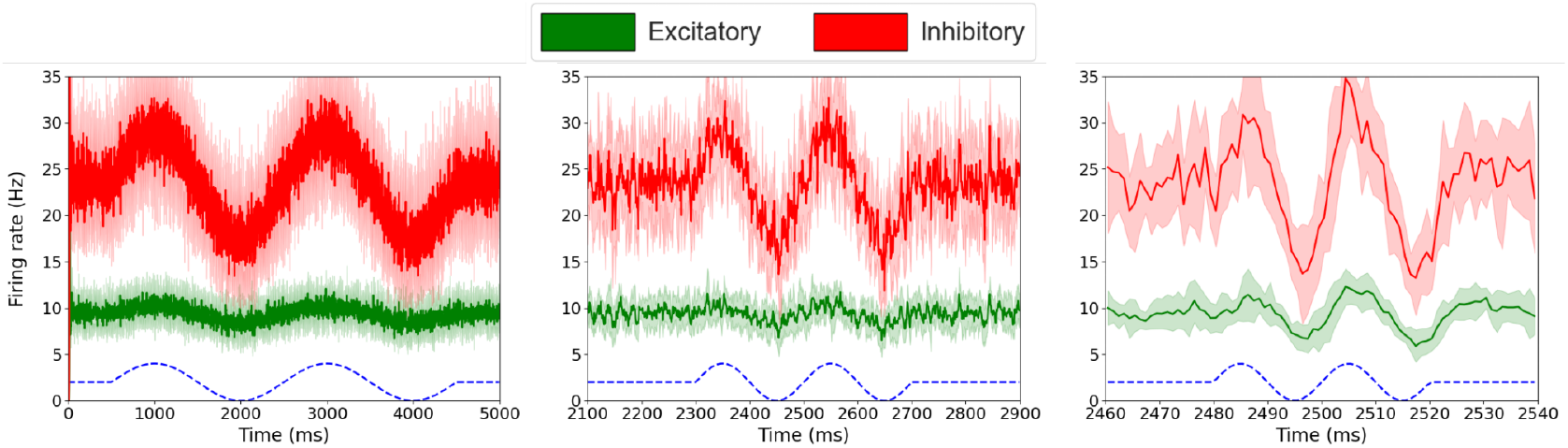
Underdamped network responses to sinusoidal inputs (two cycles). Network parameters as in Fig. 5c. **Columns 1-3:** Input frequencies equivalent to timescales in Fig. 5.

While a relatively small input amplitude (*A* = 2 Hz) was used above, we show that the peak in responsiveness to slow inputs near the working point is conserved for significantly larger input amplitudes in Supplementary Fig. S7.

### 2.5 Hodgkin-Huxley simulations

While the AdEx model reproduces many behaviours of biological neurons with only two equations, the Hodgkin-Huxley (HH) model [11] provides a more biophysically grounded description by explicitly modelling the voltage-gated ion channels underlying action potential generation. In the following, we reproduce the key result obtained from the AdEx simulations in an equivalent HH spiking network model (see 4.4). Numerical linear stabilities were not calculated for AS fixed points in the HH model, but the same analysis could be applied in principle to HH networks, and to networks composed of any neurons for which the transfer functions can be characterised [19].

Network parameters (population sizes, connection probabilities) and synaptic parameters (quantal conductances, synaptic decay constants) were unchanged from the AdEx model. Passive membrane parameters were equated to their AdEx equivalents, such that the models differ only in their spike generation mechanisms (see 4.4 for details); remaining parameters were fitted to reproduce the output firing rates of AdEx neurons at a specific operating point (*ν*_drive_ = 4 Hz, *b* = 0 pA,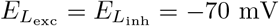).

The HH frequency-response curve (oscillatory impulses; amplitude *A* = 1 Hz) and slow-input evoked responses (*f* = 0.5 Hz) for changing 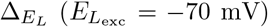 are shown in Fig. 7a and b. As in the AdEx model, the point of maximal responsiveness at the low-frequency limit coincides with the low-pass to resonant transition (at 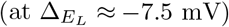. Note the presence of a second low-amplitude resonance peak at very high frequency in both network types, as observed in AdEx simulations with synaptic delays (see Supplementary Information SI.8).

**Fig. 7:**
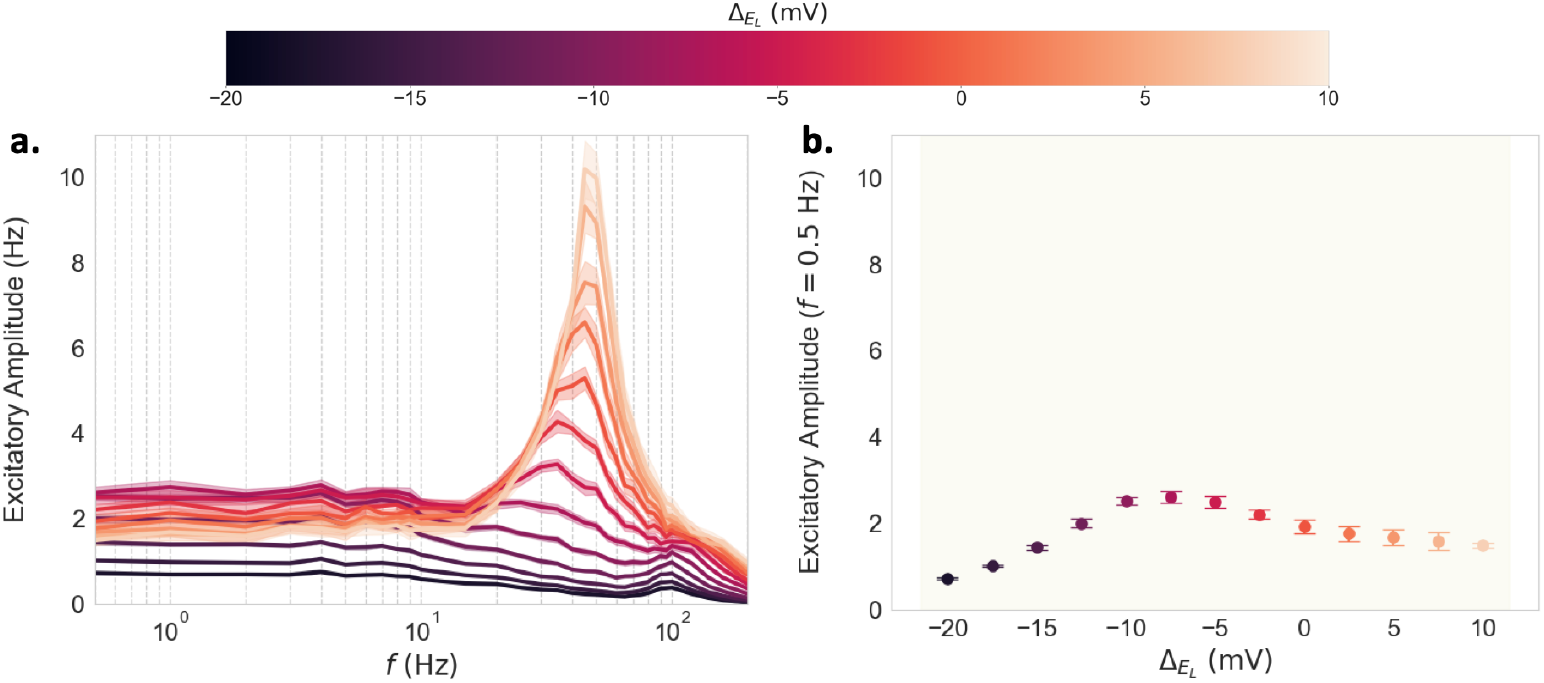
Evoked excitatory responses across 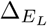 values 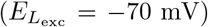 for HH simulations (*ν*_drive_ = 4 Hz, input amplitude *A* = 1 Hz). **(a)** Evoked excitatory response amplitude versus input frequency. **(b)** Evoked excitatory response amplitudes to a slow input (*f* = 0.5 Hz).

## 3 Discussion

The awake cortex faces a complex challenge: it must respond to diverse and unpredictable sensory inputs, while also retaining the capacity for frequency-selective processing when and where necessary. Here, in a cortical layer 2/3 spiking network model, we show that these dual functions are served by two distinct regimes that emerge across the continuum of asynchronous states as the excitation-inhibition balance is varied. Using a two-population semi-analytic mean-field model, we predict the responsiveness properties of the underlying high-dimensional spiking network as a function of its linear stability in a computationally efficient and generalisable manner. As the E/I balance is tuned, networks undergo a systematic transition from the broadly responsive overdamped (OD) regime to the frequency-selective underdamped (UD) regime. Crucially, across network models, control parameters and parameter regions, we find that responsiveness to slow inputs is maximised at the boundary between the two regimes at an optimal working point.

The identification of a systematic peak in responsiveness at a dynamical transition point distinguishes our approach from previous studies. Other papers have emphasised the importance of a biologically realistic conductance state [6] and have reported an inverse relationship between responsiveness and total conductance [20]. However, we show that this dependence is not monotonic: responses increase with conductance in the OD region (see Supplementary Fig. S2d). While Di Volo and Destexhe [14] also identified a peak responsiveness point in the AS regime, this maximum was a function of the heterogeneity in the network, without reference to E/I balance. As for the Ostojic [13] framework, we considered only the *homogeneous* (stable firing rate) regime; the correlates of responsiveness within this regime (in the sense explored here) were not quantitatively analysed by the author. The *heterogeneous* regime was outside the scope of this paper. Moreover, while the *afferent-dominated* (AD) and *recurrent-dominated* (RD) states identified in Zerlaut et al. [12] share features with the OD and UD regimes, respectively, the two frameworks are not directly comparable. Relative to their counterparts, the AD/OD categories show higher ratios of afferent-to-total excitatory synaptic currents (see Supplementary Fig. S2e) and correspond to more hyperpolarised, lower-conductance, lower-activity networks. However, the control parameters differ fundamentally (network-level afferent input versus population-specific intrinsic excitability), with external drive being only one factor determining the damping regime.

Whereas in Ledoux and Brunel [7], network stability was characterised as a function of connectivity parameters, here it was measured as a function of the differential excitabilities of excitatory and inhibitory neurons. Due to its direct relationship with the mean membrane potential, the leak reversal potential *E*_*L*_ was a natural parameter choice. Moreover, the largely invariant response profiles for different 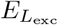 values (see Supplementary Information SI.9) was an interesting result, suggesting that the overall depolarisation level of RS and FS cells is less relevant to the network response than the difference in depolarisation between the two populations.

Given the diversity of parameters that can induce the low-pass to resonant transition *in silico*, various neuromodulatory mechanisms may be implicated in an analogous process in the awake cortex. For instance, the level of spike-dependent adaptation in RS cells (associated with AdEx parameter *b*) is inversely correlated with the concentration of acetylcholine (ACh) [18], which increases during wakefulness [21]. As well as controlling the transition between the AS and Up and Down regimes [22], the ACh-modulated level of spike-dependent adaptation may be a mechanism by which cortical regions migrate between the OD and UD regimes within the AS state. Alternatively, a neuron’s resting membrane potential (analogous to the parameter *E*_*L*_) is subject to neuromodulation by various chemicals including serotonin, which hyperpolarises and depolarises neurons via the 5-HT1A and 5-HT2A receptors, respectively [23]. In a given cortical region, if 5-HT1A and 5-HT2A receptors were to be differentially expressed in excitatory and inhibitory cells (as observed in *in vitro* experiments in layer V of the rat cortex [24]), this would effectively provide a mechanism for the serotonergic control of 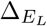.

Our results attest to the versatility of the present mean-field approach. As noted elsewhere [15], the fitting of the transfer function is performed on a single parameter set for each cell type, but its semi-analytic nature (taking various key parameters as arguments) allows it to capture the firing rates of networks with parameters very different to the fitted values. Notwithstanding, departing from the fitting region did yield a loss in accuracy. We remark that the decision to dynamically equate the mean-field timescale *T* to the autocorrelation time constant *τ*_*V*_ proved an effective choice, given the close agreement between the mean-field stability analysis and the network responses. We also stress the importance of a model with population-specific transfer functions; the slow-input maximum would not have been observed without a difference in steepness between RS and FS gains (see Supplementary Information SI.6). Moreover, this approach is generalisable to other cell types and models by fitting the transfer function to the neuron model in question (see Carlu et al. [19] for examples with the Hodgkin–Huxley and Morris–Lecar models).

We wish to emphasise the generality of these findings. The AdEx-COBA model with zero synaptic delay was analysable with an existing pre-fitted mean-field model, allowing SNN simulations to be readily complemented by the mean-field stability analysis. However, the key result in this paper – the maximum in responsiveness to slow inputs at the low-pass to resonant transition – is not merely an artefact of the AdEx model (or integrate-and-fire models in general), being replicated with the more biophysically accurate Hodgkin-Huxley model. Moreover, the result was robust to the incorporation of synaptic delays. The result also generalised beyond oscillatory inputs, extending to transient pulse-type inputs. Further, we showed that the transition between regimes is not only mediated by 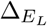, but possible with other parameters that act on the difference in excitability between RS and FS cells. Therefore, the observed transition is not exclusive to a single control parameter, but a more general phenomenon associated with the E/I balance in the network.

While the information processing implications of our findings are not explored in depth here, the systematic E/I balance-mediated transition identified at the local network scale is likely to have important functional consequences at larger spatial scales. A network in the OD regime is frequency-agnostic, responding strongly to a broad range of unknown stimuli, while UD networks act as frequency filters, with the effective frequency band depending on the amplitude of the input (see Supplementary Information SI.10). The mean-field model used here provides a natural basis for studying this dichotomy at larger scales in future work. Dynamic mean-field simulations of mesoscopic and whole-brain activity can model cortical AS states under each damping regime and compare their signal-carrying capacities; the state-dependent timescale *T* = *τ*_*V*_ can be extended naturally to such simulations at no additional computational cost (following the adaptive timescale approach of Ostojic and Brunel [25]). We hypothesise that local responsiveness and global signal transmission are correlated within the AS regime, and that networks at the working point amplify signals at the macroscopic level. From these simulated regimes, large-scale measures can be computed (including those accessible from non-invasive human brain data), such as structure-function correlation and perturbational complexity index (as in Sacha et al. [26]). These analyses may reveal macroscopic correlates of the underlying responsiveness regimes and clarify their functional roles in the awake cortex *in vivo*.

## 4 Methods

### 4.1 Spiking network model

#### 4.1.1 Single neuron model

The adaptive exponential integrate-and-fire (AdEx) model [8] was used to simulate individual neurons (equations 5 and 6):

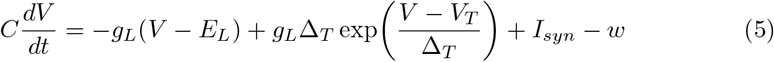

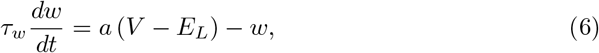

where *V* represents the neuron’s membrane potential and *w* represents a slow hyperpolarising current, due (for instance) to the action of calcium-dependent potassium ion channels [27]. When a neuron’s membrane potential reaches the spike initiation threshold *V*_*T*_, the exponential rising phase of the spike is triggered until a second threshold *V*_cut_ is reached, at which point neurons are subject to the following reset conditions:

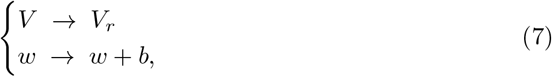

where the membrane potential is reset to a baseline value *V*_*r*_ = − 65 mV and the adaptation variable *w* is incremented by the spike adaptation current *b* pA (varied in our simulations). The membrane potential is maintained at *V*_*r*_ for a refractory time of *t*_ref_ = 5 ms after each spike. The spike adaptation current was always set to zero for FS cells (varied in RS cells), while the subthreshold (voltage-coupled) adaptation *a* was set to zero for both populations in all simulations (making *w* always zero in FS cells). Other single-cell parameters for RS and FS cells, largely taken from Sacha et al. [26] (excluding those varied), are shown in Table 1.

**Table 1:** Parameters of the AdEx single neuron model, COBA synaptic model and spiking network model for RS and FS cells.

| Description | Symbol | Cell type |  | Units |
| --- | --- | --- | --- | --- |
|  |  | RS | FS |  |
| Membrane capacitance | $C_m$ | 200 | 200 | pF |
| Leak conductance | $g_L$ | 10 | 10 | nS |
| Leak reversal potential | $E_L$ | -70 | varied | mV |
| Spike initiation threshold | $V_T$ | -50 | -50 | mV |
| Slope factor | $\Delta_T$ | 2 | 0.5 | mV |
| Spike cut-off threshold | $V_{\text{cut}}$ | -40 | -47.5 | mV |
| Reset potential | $V_r$ | -65 | -65 | mV |
| Refractory period | $t_{\text{ref}}$ | 5.0 | 5.0 | ms |
| Adaptation time constant | $\tau_w$ | 500 | - | ms |
| Subthreshold adaptation parameter | $a$ | 0 | - | nS |
| Spike-triggered adaptation current | $b$ | varied | - | pA |
| Excitatory synaptic reversal potential | $E_e$ | 0 | 0 | mV |
| Inhibitory synaptic reversal potential | $E_i$ | -80 | -80 | mV |
| Excitatory quantal conductance | $Q_e$ | 1.5 | 1.5 | nS |
| Inhibitory quantal conductance | $Q_i$ | 5.0 | 5.0 | nS |
| Excitatory synaptic decay constant | $\tau_e$ | 5.0 | 5.0 | ms |
| Inhibitory synaptic decay constant | $\tau_i$ | 5.0 | 5.0 | ms |
| Network population size | $N_e / N_i$ | 8000 | 2000 | - |
| Probability of connection<br>(between and within populations) | $p$ | 5% | 5% | - |
| External drive rate | $\nu_{\text{drive}}$ | varied | varied | Hz |

#### 4.1.2 Synaptic model

The synaptic model is summarised in equations 8 and 9 below. Instantaneous conductance-based (COBA) synapses with exponential decay and zero delay were used (the effect of synaptic delays is briefly explored in Supplementary Information SI.8).

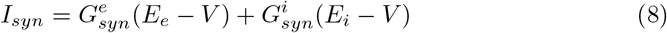

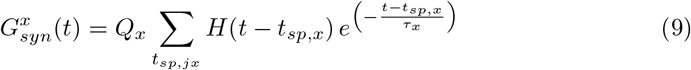

Here, *H* represents the Heaviside step function:

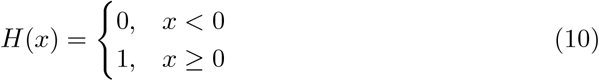

For a given neuron, its excitatory (inhibitory) conductance 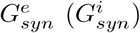 mented by quantal conductance *Q*_*e*_ (*Q*_*i*_) for each excitatory (inhibitory) presynaptic spike it receives at time *t*_*sp,e*_ (*t*_*sp,i*_). Parameters are shown in Table 1.

#### 4.1.3 Layer 2/3 cortical network model

Networks were constructed by simulating a population of 10000 neurons with sparse connectivity (5% probability of connection between all neurons, excluding self-connections), divided into a population of *N*_*e*_ = 8000 RS cells and *N*_*i*_ = 2000 FS cells. An excitatory drive from an external population of *N*_*e*_ = 8000 excitatory Poissonian neurons with mean firing rate *ν*_drive_ was applied to both populations (with 5% connection probability) to simulate background noise (see Fig. 1a). The Brian 2 simulator [28] was used for all spiking network simulations.

### 4.2 Measuring responsiveness

Network responsiveness was measured by the amplitude of the evoked population responses to time-varying sinusoidal drives, computed by fitting sinusoids (least squares) to population firing rates (Fig. 1b) and averaging over 10 seeds. All other SNN metrics in this paper were also averaged over 10 seeds, with error bars indicating standard deviations in all plots.

The afferent sinusoidal drive (approximating a rhythmic input from the thalamus or a neighbouring cortical region) was generated by varying the mean firing rate of the external population of 8000 excitatory Poisson neurons about the baseline drive rate, *ν*_drive_, of the network in question.

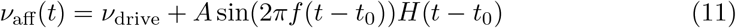

The oscillation was initiated after a baseline period of *t*_0_ = 2000 ms to ensure the passage of the transient at the simulation onset, and the fit was performed on the last 2000 ms of the simulation. The frequency *f* and amplitude *A* of the afferent oscillation were varied across simulations.

### 4.3 Linear stability analysis: a mean-field approach

The study of the linear stability of nonlinear integrate-and-fire networks with COBA synapses does not admit of a simple analytical approach. To this end, an adaptive mean-field model derived from the spiking network was used to numerically compute the linear stabilities of a range of stable AS fixed points as the E/I balance of the network was varied. This method provided an efficient means of numerically approximating the stability of a large number of AS fixed points without the need to estimate the eigenvalues from SNN simulations (wherein the spiking network is perturbed and the partial derivatives are estimated directly), which is time-consuming and subject to large trial-by-trial variance.

The difference between excitatory and inhibitory leak potentials, 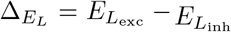, a proxy for the E/I balance of the network, was chosen as a control parameter, but the results generalise to other parameters playing on this balance (see Supplementary Information SI.4). For a given 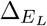, population firing rates and synaptic conductances are highly conserved when 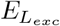 and 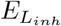 are varied over a broad physiologically-relevant range. Consequently, their eigenvalues and responsiveness profiles are largely invariant (see Supplementary Information SI.9), a possible source of degeneracy for the modulation of cortical responses under different neuromodulatory conditions. Unless otherwise stated, we fixed 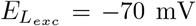 and varied 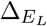 via changes in 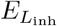. Eigenvalues of the linearised two-dimensional system (fixing adaptation at its stationary value) were computed, yielding either negative real eigenvalues (stable nodes) or complex conjugate eigenvalues (stable foci). Details of the mean-field model and linear stability calculations are given below.

#### 4.3.1 Mean-field equations

Linear stability calculations were performed using the first-order reduction of the mean-field model with adaptation introduced in Di Volo et al. [15]. The threedimensional system of equations is shown below, where *ν*_*e*_ (*ν*_*i*_) represents the mean excitatory (inhibitory) population firing rate, *w* represents the mean adaptation current for excitatory neurons (with spike adaptation current *b* equal to its SNN analogue and subthreshold adaptation *a* set to 0), and *F*_*µ*_ is a semi-analytic transfer function (input-output function) fitted to neurons in population *µ* ∈ (*e, i*):

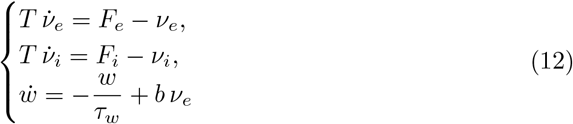

The population rate equations are identical to the Wilson-Cowan model with the exception of the transfer function used, the details of which are given below.

#### 4.3.2 Neuronal transfer function

The transfer functions *F*_*µ*_(*ν*_*e*_, *ν*_*i*_, *w*) (that is, the functions relating a neuron’s output firing rate to its excitatory and inhibitory presynaptic release rates, *ν*_*e*_ and *ν*_*i*_, and the adaptation current *w*) were derived semi-analytically based on the approach introduced in Zerlaut et al. [17] and adapted to account for adaptation in Di Volo et al. [15].

The instantaneous output firing rate, 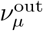, of a single neuron in population *µ* can be expressed as a function of its membrane potential fluctuation statistics (mean *µ*_*V*_, standard deviation *σ*_*V*_ and autocorrelation time constant *τ*_*V*_) at a given point in time, as follows:

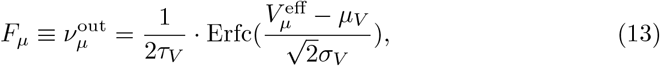

where Erfc is the complementary error function and 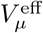 is a phenomenological spiking threshold, fitted separately to each neuron type *µ* as a second-order polynomial combination of the subthreshold moments. The polynomial fits performed on RS and FS cells from Di Volo et al. [15] were used in this study.

The subthreshold moments are obtained from the instantaneous excitatory and inhibitory presynaptic input rates, *ν*_*e*_ and *ν*_*i*_, following Campbell’s Theorem, on the assumption that the synaptic inputs follow a Poisson distribution [29, 30], which is approximately true for networks in the AS firing regime. Details of the derivations of the subthreshold moments and fitted polynomial coefficients can be found in Supplementary Information SI.11.

#### 4.3.3 Computing eigenvalues

The jacobian matrix *J* ^∗^ ∈ ℝ^2×2^ of the linearised two-dimensional system at fixed point 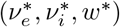 (fixing the adaptation current at its stationary value, *w*^∗^) is given below:

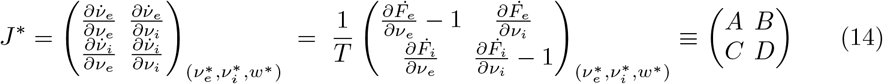

The characteristic equation of *J* ^∗^ is therefore

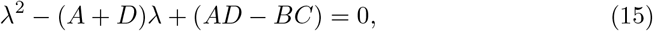

yielding the eigenvalues:

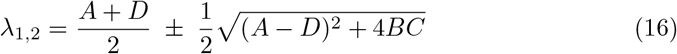

Partial derivatives were computed numerically using the central differences method with an adaptive step size as follows:

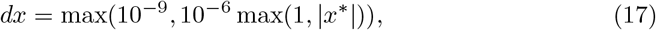

where *x*^∗^ is the equilibrium value of *x* ∈ {*ν*_*e*_, *ν*_*i*_}.

#### 4.3.4 State-dependent mean-field timescale *T* = *τ*_*V*_

The eigenvalues scale inversely with the mean-field timescale, *T*, making its choice critical to the accuracy of the stability analysis. The choice of mean-field timescale *T* has been discussed at length in various papers [15, 31, 32]. Theoretically, the optimal value for this parameter is the smallest interval for which the assumption of Markovian dynamics applies to the network [31], that is, the minimal *T* for which the network depends only on its state after time *t* − *T* and is independent of its state at all earlier times.

The mean global autocorrelation of the membrane potential fluctuations, *τ*_*V*_, is the theoretical lower bound on this timescale and depends dynamically on the network state; a single choice of *T* fails to capture the variance in the network autocorrelation times for different activity levels. To this end, in all calculations performed here, the state-dependent timescale *T* = *τ*_*V*_ has been used, with the stationary value of *τ*_*V*_ extracted from the excitatory transfer function for a given fixed point (inhibitory *τ*_*V*_ values were identical to excitatory values to very high precision). The *T* values corresponding to each of the networks explored in the Results section are shown in Fig. 8.

**Fig. 8:**
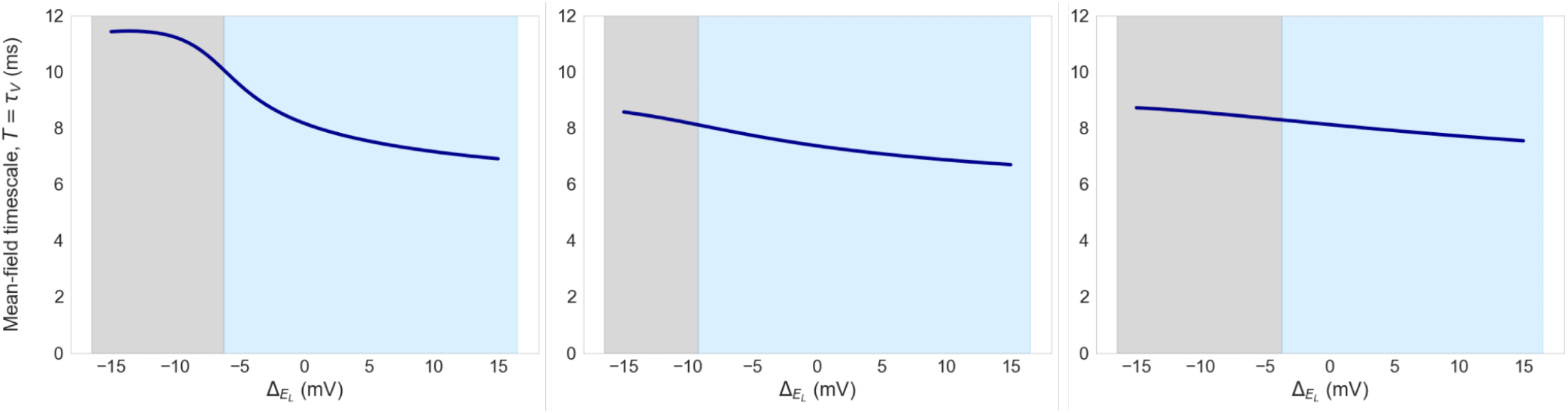
State-dependent mean-field timescale *T* versus 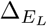, equated to the stationary value of the autocorrelation time constant *τ*_*V*_ from the excitatory neuronal transfer function for the associated 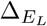 value. Grey (blue) represents the overdamped (underdamped) region. **Columns 1-3**: parameter sets as in Fig. 2.

### 4.4 Hodgkin-Huxley model

#### 4.4.1 Model equations

The five-dimensional system of equations defining the Hodgkin-Huxley model is shown below [33]:

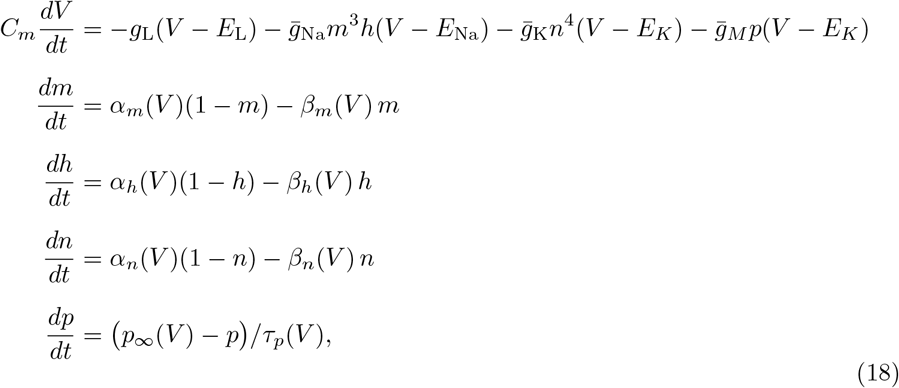

where the 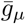 terms represent the maximal conductances for the sodium (Na) current, the delayed-rectifier potassium (K) current, and the slow non-inactivating potassium M-current (M); *E*_Na_ and *E*_*K*_ are the corresponding reversal potentials, while *g*_L_, *E*_L_ and *C*_*m*_ are identical to their AdEx equivalents. The gating variables *m* and *h* respectively describe the activation and inactivation of the Na channel, *n* describes the activation of the delayed-rectifier K^+^ channel, and *p* describes the activation of the M-current channel. Each *α*_*µ*_(*V*) and *β*_*µ*_(*V*) is a voltage-dependent rate constant governing the opening and closing of the corresponding gate, while *p*_∞_(*V*) and *τ*_*p*_(*V*) are the steady-state activation and voltage-dependent time constant of the M-current gate.

The voltage-dependent rate constants and steady-state gating functions are given by:

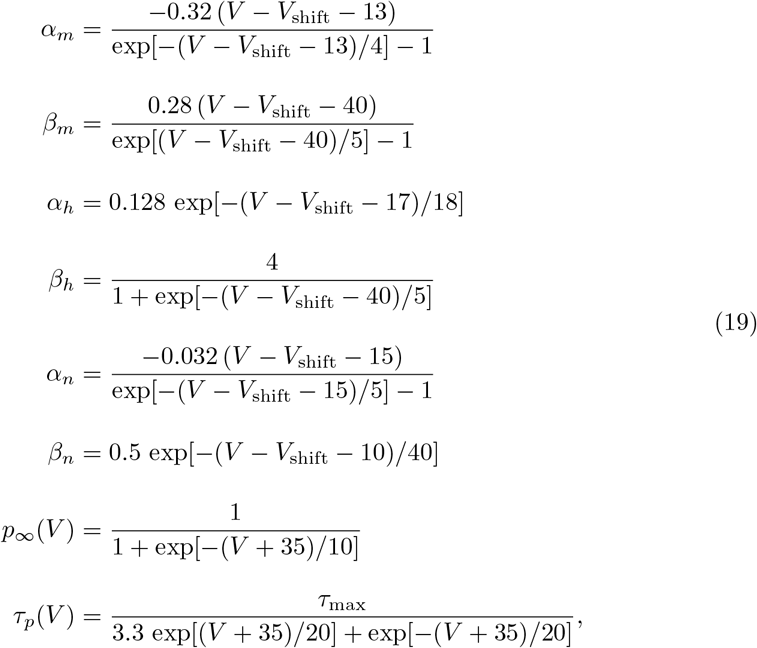

where *V*_shift_ is the voltage threshold shift parameter, modulating the spike activation and inactivation ranges, and *τ*_max_ is the maximal value of the M-current time constant *τ*_*p*_(*V*).

#### 4.4.2 Parameterisation

The network architecture, connection probabilities, and synaptic parameters are unchanged from AdEx simulations (see 4.1.2 and 4.1.3). To generate comparable parameter sets to the AdEx network, the passive membrane parameters in the HH cells were made identical to the associated AdEx values: the leak conductance (*g*_*L*_) and membrane capacitance (*C*_*m*_) were equated to 10 nS and 200 pF, respectively, in both cell types. To remove adaptation effects, the M-current maximum conductance 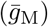 was equated to zero. With initial estimates based on experimental recordings from Pospischil et al. [33], a grid search was then performed to optimise the maximum sodium conductances 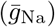, maximum potassium conductances 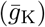 and voltage thresholds (*V*_shift_) of RS and FS cells to match the firing rates of corresponding AdEx cells (at an operating point of: *ν*_drive_ = 4 Hz, *b* = 0 pA,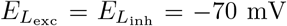). Example membrane potential traces of AdEx and HH neurons (under the predicted level of synaptic bombardment at the operating point) are shown in Fig. 9.

**Fig. 9:**
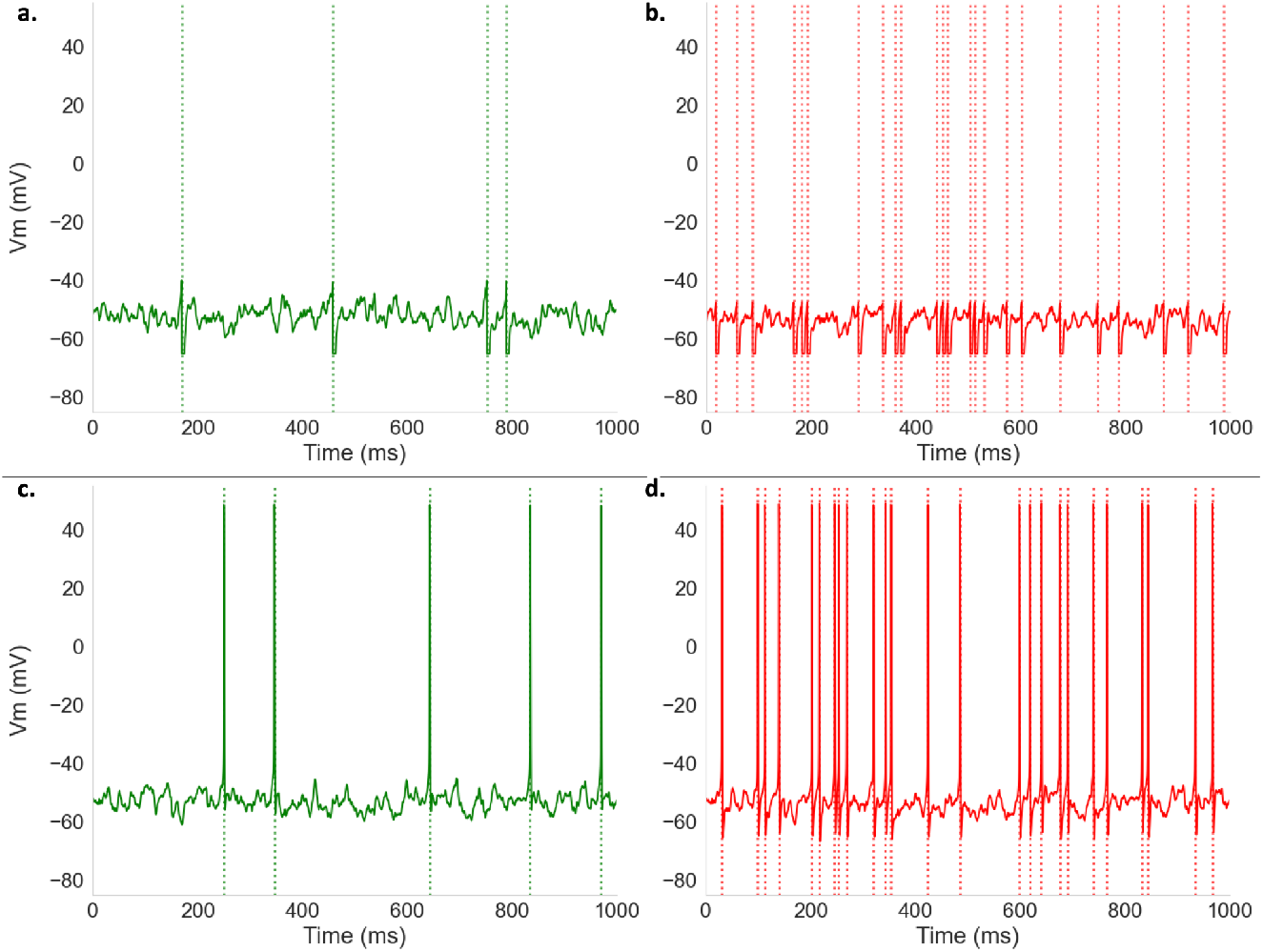
Sample single neuron membrane potential traces for AdEx and HH neurons. **(a)** AdEx, RS. **(b)** AdEx, FS. **(c)** HH, RS. **(d)** HH, FS.

The parameter sets resulting from the grid search are listed in Table 2.

**Table 2:** Single-cell parameters of the Hodgkin-Huxley model for RS and FS cells.

| Description | Symbol | Cell type |  | Units |
| --- | --- | --- | --- | --- |
|  |  | RS | FS |  |
| Membrane area | $A$ | 200 | 200 | $\mu\text{m}^2$ |
| Membrane capacitance | $C_m$ | 1.0<sup>1</sup> | 1.0<sup>1</sup> | $\mu\text{F}/\text{cm}^2$ |
| Leak conductance | $g_L$ | 0.05<sup>2</sup> | 0.05<sup>2</sup> | $\text{mS}/\text{cm}^2$ |
| Leak reversal potential | $E_L$ | -70 | -70 | mV |
| Na<sup>+</sup> max conductance | $\bar{g}_{\text{Na}}$ | 90<sup>3</sup> | 110<sup>4</sup> | $\text{mS}/\text{cm}^2$ |
| K<sup>+</sup> max conductance | $\bar{g}_{\text{K}}$ | 3.0<sup>5</sup> | 5.0<sup>6</sup> | $\text{mS}/\text{cm}^2$ |
| M-current max conductance | $\bar{g}_M$ | — | — | $\text{mS}/\text{cm}^2$ |
| Na<sup>+</sup> reversal potential | $E_{\text{Na}}$ | 50 | 50 | mV |
| K<sup>+</sup> reversal potential | $E_{\text{K}}$ | -90 | -90 | mV |
| Voltage shift | $V_{\text{shift}}$ | -53 | -55 | mV |
| M-current time constant | $\tau_{\text{max}}$ | — | — | ms |

<sup>1</sup> 200 pF; <sup>2</sup> 10 nS; <sup>3</sup> 18,000 nS; <sup>4</sup> 22,000 nS; <sup>5</sup> 600 nS; <sup>6</sup> 1,000 nS

## Acknowledgements

Research supported by CNRS, the European Union (Virtual Brain Twin project, Horizon Health 101137289), and Agence Nationale de la Recherche (FLAG-ERA grant BrainAct, CR-CNS grant ImpactCom).

## Code availability

The code necessary to reproduce all figures in the main article is available at https://github.com/mattbassat/async responsiveness. Code for Hodgkin-Huxley simulations was adapted from Carlu et al. [19]. Additional code is available upon request.

## Supplementary Information

### SI.1 Evoked responses to larger-amplitude oscillatory impulses (*A* = 2 Hz)

In Fig. S1, we show the equivalent results to those shown in Fig. 2 for oscillatory impulses with amplitude *A* = 2 Hz.

**Fig. S1:**
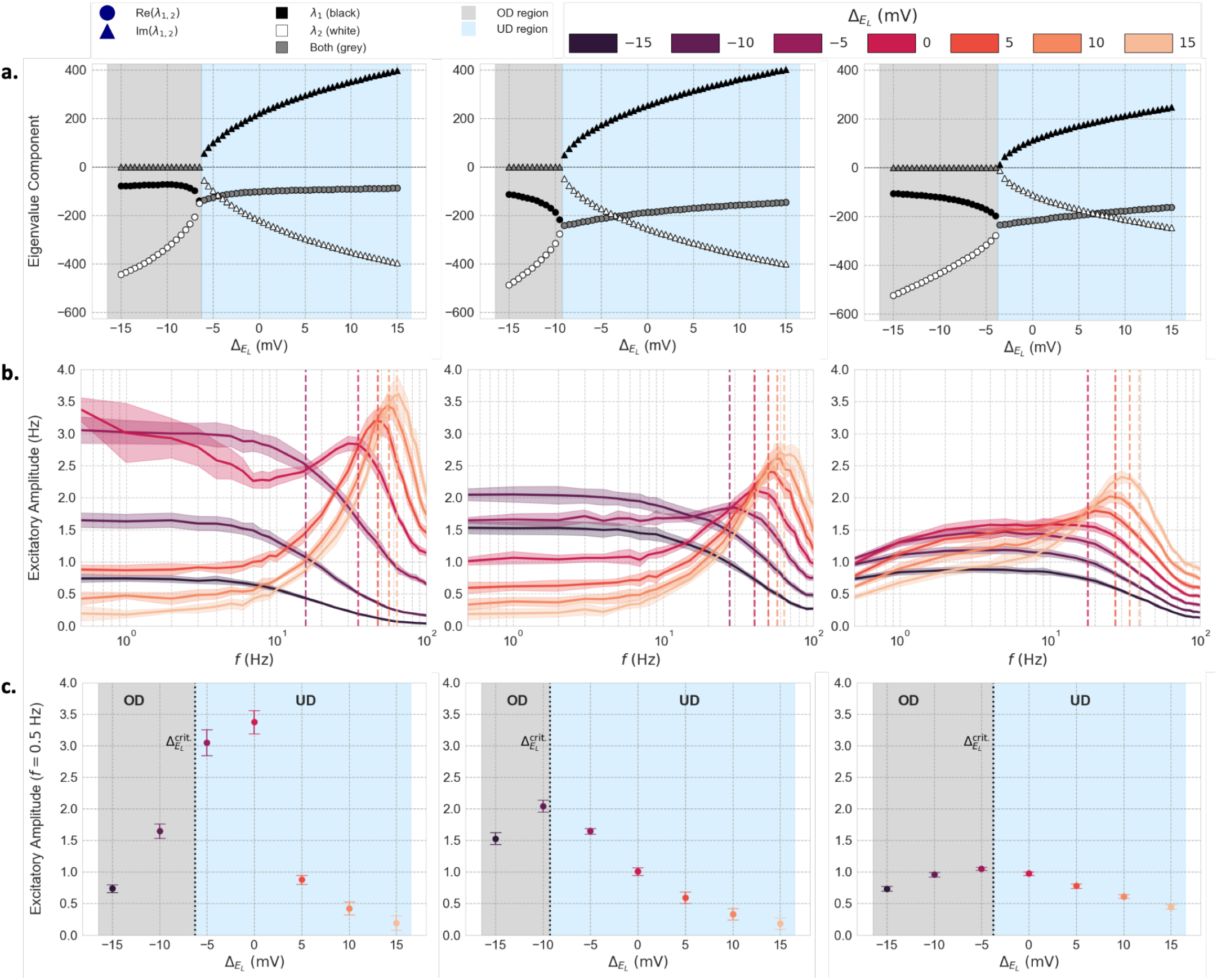
Linear stability analysis and frequency-response profiles across parameter combinations (input amplitude *A* = 2 Hz). Columns and rows as in Fig. 2.

### SI.2 Baseline activity metrics

Below, we report various baseline metrics for the networks shown in Fig. 2. All values are averaged over 10 seeds with a simulation time of 5000 ms (ignoring an initial transient period of 500 ms).

In Fig. S2d, the conductance ratio 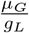 is equal to the average total conductance divided by the leak conductance, that is, 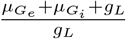.

In Fig. S2e, the baseline afferent-to-total excitatory synaptic current ratio (grey) is computed as follows (by separating the synaptic currents into those from the external and internal excitatory populations):

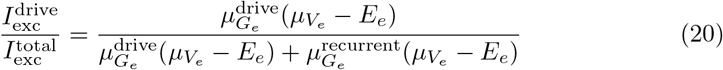

The inhibitory-to-excitatory synaptic current ratio is computed as follows:

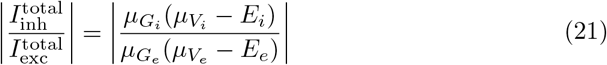

**Fig. S2:**
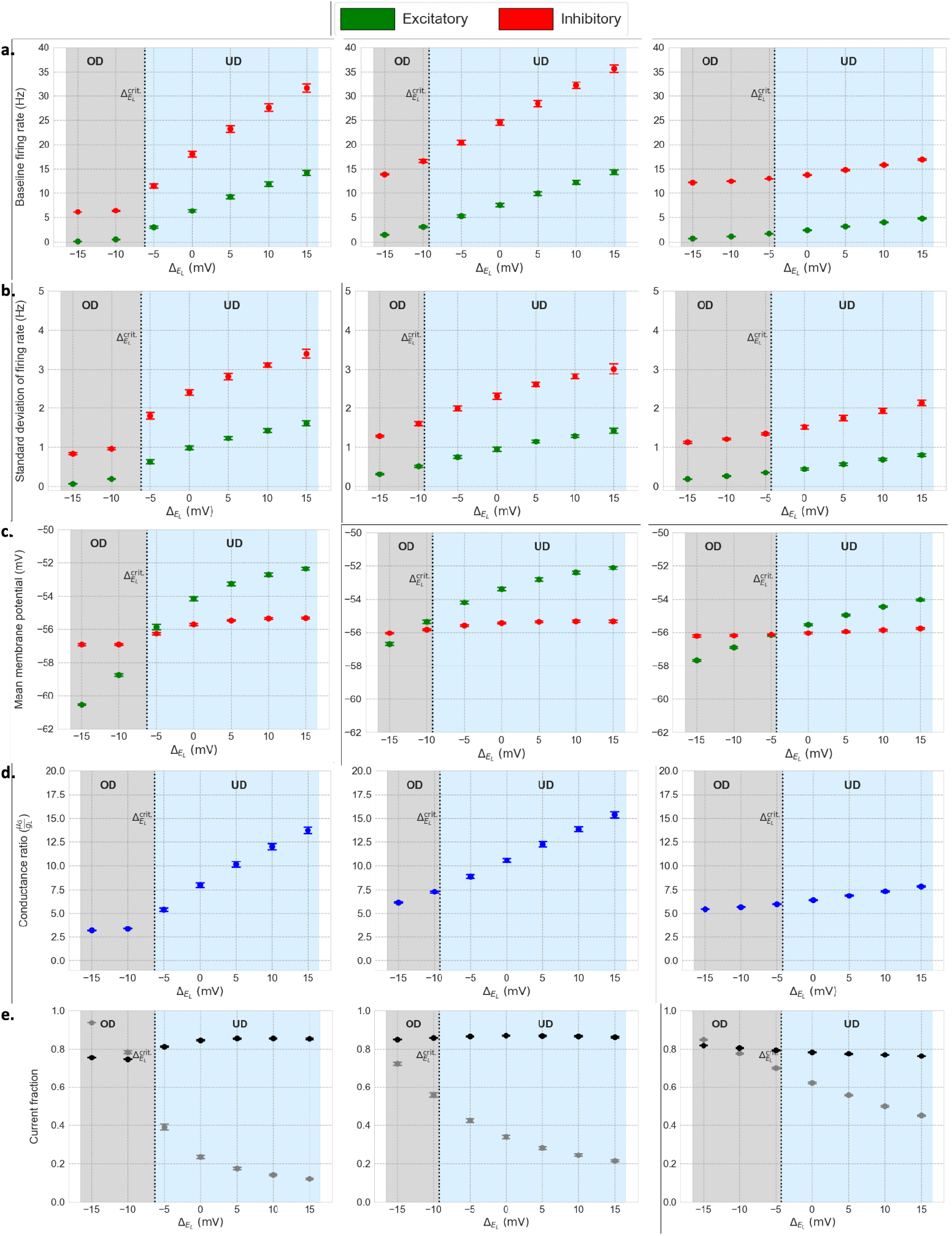
Baseline excitatory and inhibitory population statistics for networks in Fig. 2. **(a)** Mean firing rate. **(b)** Standard deviation of firing rate. **(c)** Average membrane potential. **(d)** Average conductance ratio, 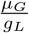. **(e)** Ratio of afferent excitatory synaptic current to total excitatory synaptic current, 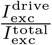 (grey) and absolute ratio of inhibitory synaptic current to excitatory synaptic current, 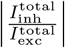 (black), as in Zerlaut et al. [12]. **Columns 1-3:** parameter sets as in Fig. 2.

### SI.3 Derivation of analytical expression for evoked responses to slow inputs

After a small perturbation in the afferent drive from its equilibrium value, *δν*_drive_, engendering small perturbations *δν*_*e*_ and *δν*_*i*_ in the excitatory and inhibitory firing rates, we write the first-order Taylor expansions of the stationary transfer functions *F*_*e*_ and *F*_*i*_ as follows:

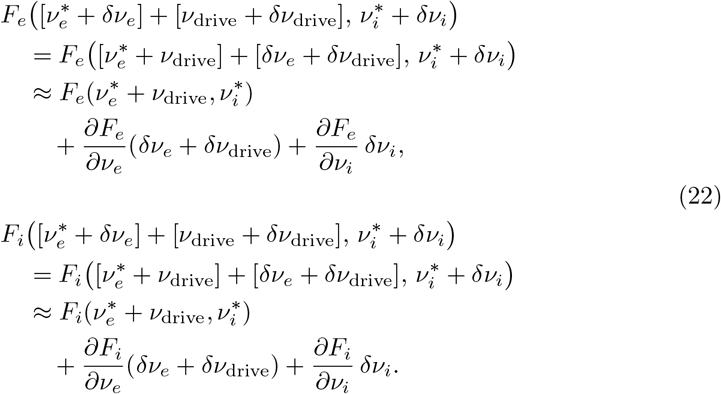

yielding the following expressions for the dynamics (from 12):

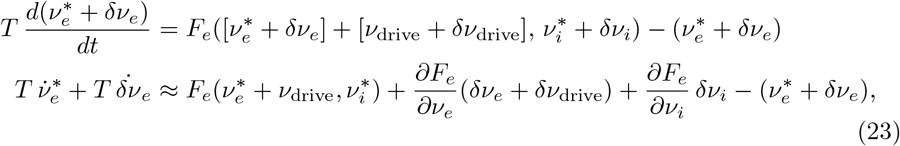

and analogously for inhibition. From equation 12, this simplifies to:

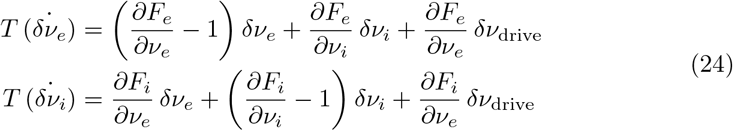

Expressing 24 in matrix form, we obtain:

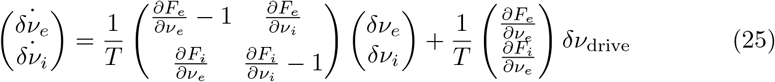

Equivalently:

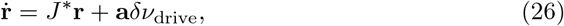

where **r** = (*δν*_*e*_, *δν*_*i*_)^⊤^, *J* ^∗^∈ ℝ^2×2^ is the jacobian matrix (4.3.3) and **a** is the vector 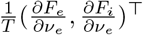 evaluated at the fixed point in question.

### SI.4 Generalisation of damping transition to other cellular parameters

The same qualitative transition in linear stability (and, by extension, responsiveness mode) is observed for other parameters that modulate the differential excitability of the inhibitory and excitatory populations. As in the case of 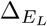, maximal slow-input responsiveness consistently coincides with the low-pass to resonant boundary.

In Fig. S3, we fix the leak potentials 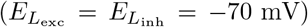 and vary the difference between population leak conductances, 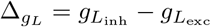.

**Fig. S3:**
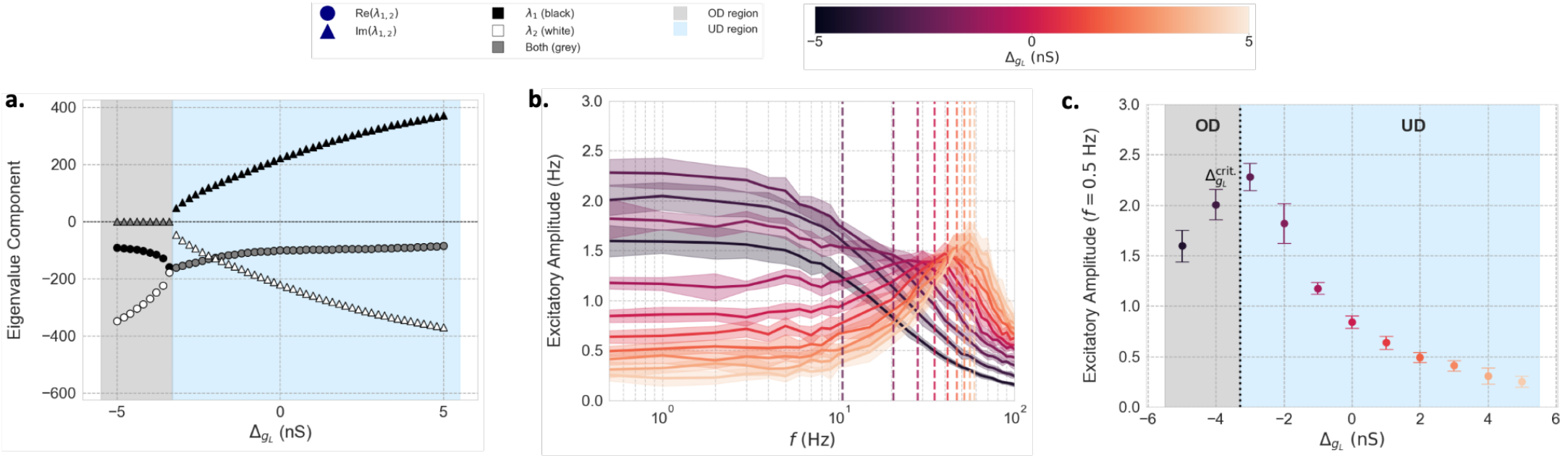
Mean-field linear stability analysis and SNN frequency-response profiles (input amplitude *A* = 1 Hz) for changing 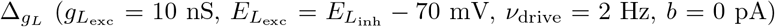. **(a)** Eigenvalues versus 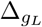. Shapes indicate real (circles) and imaginary (triangles) parts. Colours denote eigenvalue index (black: *λ*_1_; white: *λ*_2_; dark grey: common to both). Background colours denote OD (light grey) and UD (blue) regimes. **(b)** Amplitudes of evoked excitatory responses versus input frequency *f* ∈ [0.5, 100] Hz (log scale), coloured by 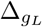. Dashed lines: eigenfrequencies for UD subset. **(c)** Visualisation of evoked responses to a slow input (*f* = 0.5 Hz). 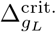 denotes the mean-field prediction of the boundary between OD (grey) and UD (blue) regions.

In Fig. S4, we vary *b*, the spike-adaptation current in RS neurons. Note that the mean-field linear stability analysis is highly inaccurate for large *b* (given the transfer function was fitted at a value of *b* = 0 pA). The position of the resonance peak is well captured for *b* = 0 pA, but the value of the working point *b*^crit.^ is significantly overestimated, with the true value (marking the transition from resonant to nonresonant) occurring at *b* ≈ 25 pA (all networks from *b* = 50 pA to *b* = 100 pA were therefore misclassified as UD networks). Notwithstanding, the same general result was obtained, that is, the occurrence of the slow-input maximum near the transition point between OD and UD regimes.

**Fig. S4:**
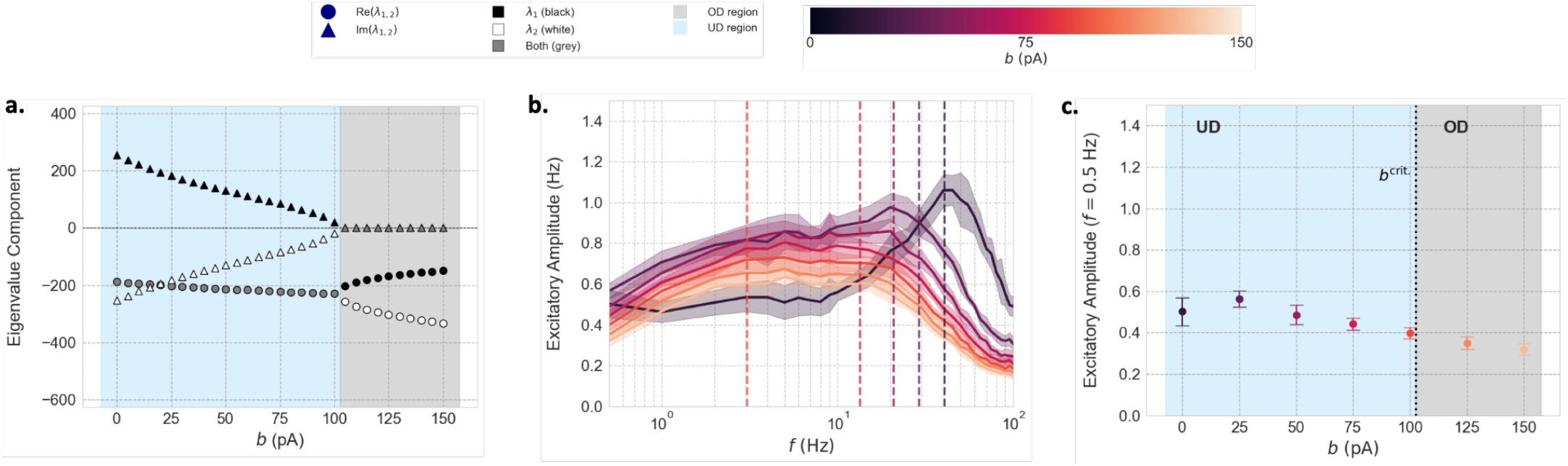
Mean-field linear stability analysis and SNN frequency-response profiles (input amplitude *A* = 1 Hz) for changing 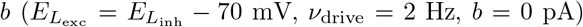. **(a)** Eigenvalues versus *b*. Shapes indicate real (circles) and imaginary (triangles) parts. Colours denote eigenvalue index (black: *λ*_1_; white: *λ*_2_; dark grey: common to both). Background colours denote OD (light grey) and UD (blue) regimes. **(b)** Amplitudes of evoked excitatory responses versus input frequency *f* ∈ [0.5, 100] Hz (log scale), coloured by 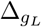. Dashed lines: eigenfrequencies for UD subset. **(c)** Visualisation of evoked responses to a slow input (*f* = 0.5 Hz). 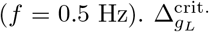 denotes the mean-field prediction of the boundary between OD (grey) and UD (blue) regions.

### SI.5 Responsiveness results for higher external drive (*ν*_drive_ = 8 Hz)

Responsiveness results (*A* = 1 Hz) and mean-field linear stability calculations for *ν*_drive_ = 8 Hz and *b* = 0 pA are shown in Fig. S5. Here, as in the high-adaptation (*b* = 60 pA) case, the transfer function departs significantly from its fitting range and, consequently, the mean-field linear stability calculations are less accurate; while the mean-field estimated a working point of 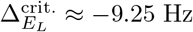, the true value (as revealed in subsequent SNN simulations) is between 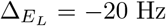 and 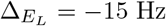. The positions of the resonance peaks (i.e. |Im{*λ*_1,2_}|) are also significantly underestimated. The same qualitative changes in responsiveness are seen from *ν*_drive_ = 4 Hz to *ν*_drive_ = 8 Hz as from *ν*_drive_ = 2 Hz to *ν*_drive_ = 4 Hz: a decrease in the (true) working point, a relatively small increase in the resonant frequencies, a large reduction in the heights of the resonance peaks, a reduction in the height of the responsiveness peak at the damping boundary, and a global flattening of the responses across the parameter range and across frequencies.

**Fig. S5:**
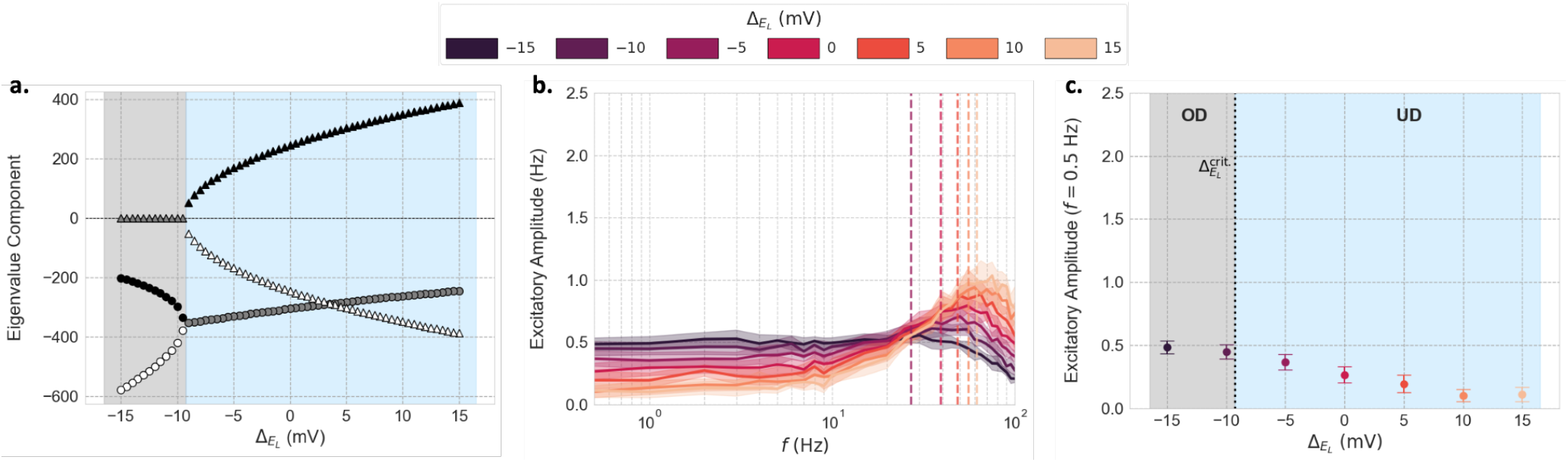
Mean-field linear stability analysis and SNN frequency-response profiles (*A* = 1 Hz) versus 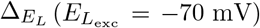 for *ν*_drive_ = 8 Hz. **(a)** Eigenvalues versus 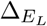. **(b)** Amplitudes of evoked excitatory responses versus input frequency, coloured by 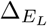. Dashed lines: eigenfrequencies for UD subset. **(c)** Visualisation of evoked responses to a slow input (*f* = 0.5 Hz).

### SI.6 Slow-input responsiveness: gain analysis

Per Eq. 4, the evoked perturbations 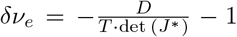 and 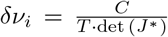 are inversely proportional to det (*J* ^∗^) = *AD* − *BC* = |*BC*| − |*AD*|, which is minimised in the vicinity of the damping boundary, as shown in Fig. S6a.

This can be framed as a competition between the ‘self gain’ and ‘coupling’ terms. The product of self gains |*AD*| drives the system towards excitation, since *A* promotes an increase in *ν*_*e*_ and *D* promotes a decrease in *ν*_*i*_. Conversely, the product of coupling gains |*BC*| has a dampening effect on the system, since *B* promotes a reduction in *ν*_*e*_ and a *C* promotes an increase in *ν*_*i*_. The determinant is minimised when |*AD*| is maximised relative to |*BC*|, that is, when the excitability of the system due to the E-E and I-I loops is largest relative to the inhibiting influence of the E-I coupling.

In the OD region, the RS population is in a low activity regime wherein the excitatory gains *A* and *B* are small relative to the inhibitory gains *C* and *D*, since FS cells are significantly closer to threshold. As 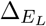 increases, the RS cells enter the steep rising phase of their transfer functions, yielding a faster growth in |*AD*| than in |*BC*| due to the proportionately larger increase in the E-E gain *A* than in E-I gain *B*, and a consequent decrease in det (*J* ^∗^).

As 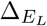 approaches 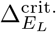 from the left, the rates of change in the excitatory gains slow down and reach an inflexion point (Fig. S6b). Here, det (*J* ^∗^) is minimised and the gains are at a balancing point, that is, 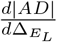 and 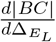 are equal (Fig. S6c):

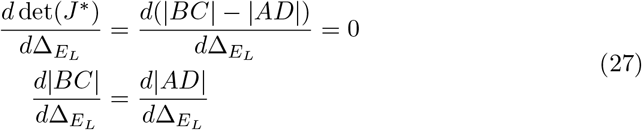

Beyond the damping boundary, 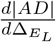 becomes less positive than 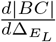 due to the disproportionately large decrease in 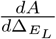, reflecting the rapid saturation of RS self-excitation relative to E–I coupling. This yields a slower increase in |*AD*| than in |*BC*| and an attendant increase in det (*J* ^∗^).

**Fig. S6:**
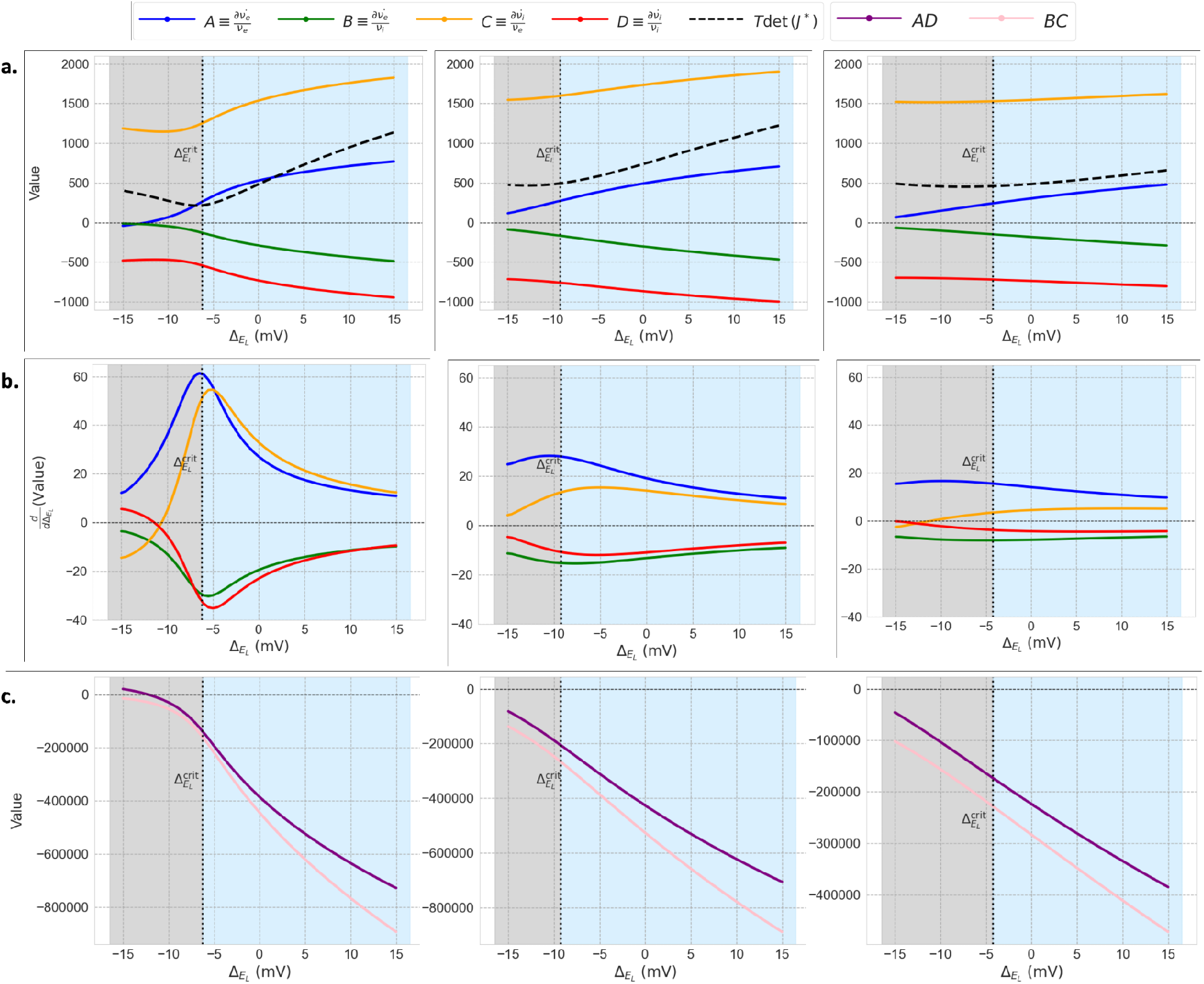
Maximal slow-input responsiveness at 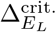 coincides with a minimum in the determinant det (*J* ^∗^), where the ‘self gain’ and ‘coupling gain’ terms have equal gradient and the difference between them is minimised. **(a)** Mean-field partial derivatives and determinants of the jacobian matrices of AS fixed points across 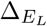 values. **(b)** Rates of change with respect to 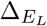 of partial derivatives. **(c)** Self gain (*AD*) and coupling (*BD*) gain terms as a function of 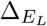, **Columns 1-3**: parameter sets as in Fig. 2.

### SI.7 Slow-input responses to high-amplitude pulse inputs

In Fig. S7, we show that the slow-input responsiveness peak generalises to pulse inputs (as in 2.4) of arbitrarily high amplitude. Responses to sinusoidal half-cycle pulses of duration 1000 ms (as in Fig. 5, column 1) were computed across the 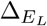 range for the networks with parameters *ν*_drive_ = 2 Hz, *b* = 0 pA. Evoked amplitudes were calculated by an identical fitting procedure to that used for oscillatory impulses (see 4.2), but this time over the half-cycle stimulus period.

**Fig. S7:**
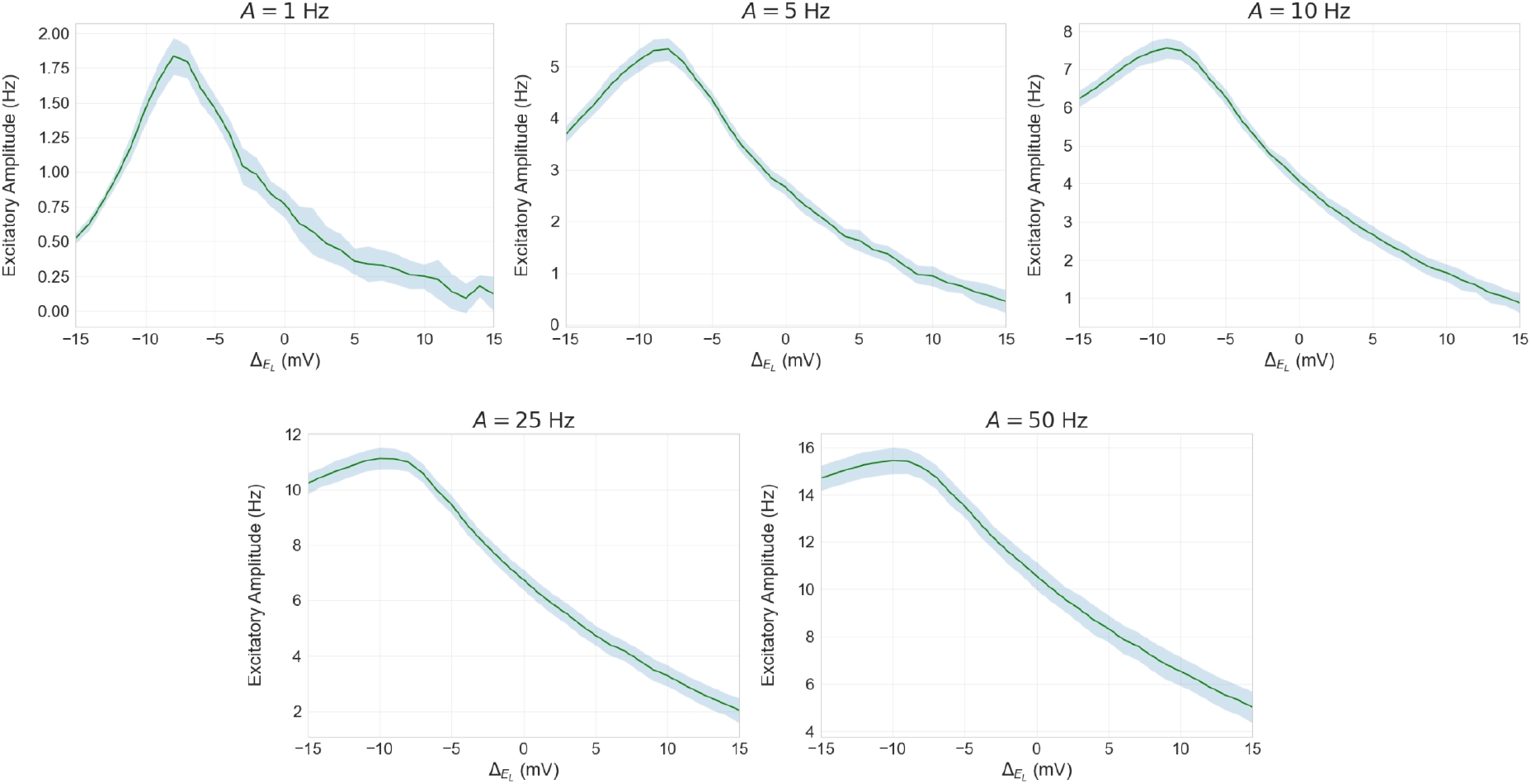
Evoked excitatory responses to a slow pulse input (duration 1000ms, equivalent to an oscillatory frequency of *f* = 0.5 Hz) versus 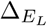 for various input amplitudes *A* (*ν*_drive_ = 2 Hz, *b* = 0 pA).

### SI.8 Evoked responses with non-zero synaptic delays

In Fig. S8, we re-run the responsiveness simulations for *ν*_drive_ = 4 Hz and *b* = 0 pA with the addition of a small uniform synaptic delay in excitatory and inhibitory synapses (first column: 1 ms; second column: 2 ms).

The incorporation of synaptic delays has a significant effect on network stability. Firstly, networks become increasingly unstable with increasing delay, as evidenced by the growth in resonance peak heights as the delay increases from zero (Fig. 2b, second column) to 1 ms, and from 1 ms to 2 ms. The networks with the highest 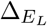 values (i.e. the most unstable networks) undergo Hopf bifurcations and exit the AS regime for sufficiently large delay, hence the exclusion of 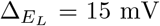 from column 1 and the exclusion of 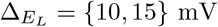 from column 2. Secondly, the delays introduce a second, lower-amplitude resonance peak at very high frequencies (*>* 100 Hz) due to inhibitory-inhibitory feedback [7]. This high-frequency peak appears in both resonant and low-pass networks.

Despite these effects, the maximal responsiveness to slow inputs still coincides with the transition point between the low-pass and resonant regimes in the presence of delays (Fig. S8b). Moreover, in all three cases (zero delay, delay=1 ms and delay=2 ms), the critical 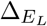 value is unchanged, and the slow-input evoked responses are almost identical. Therefore, the maximal slow-input responsiveness at the damping boundary is highly robust to the inclusion of realistic synaptic delays.

**Fig. S8:**
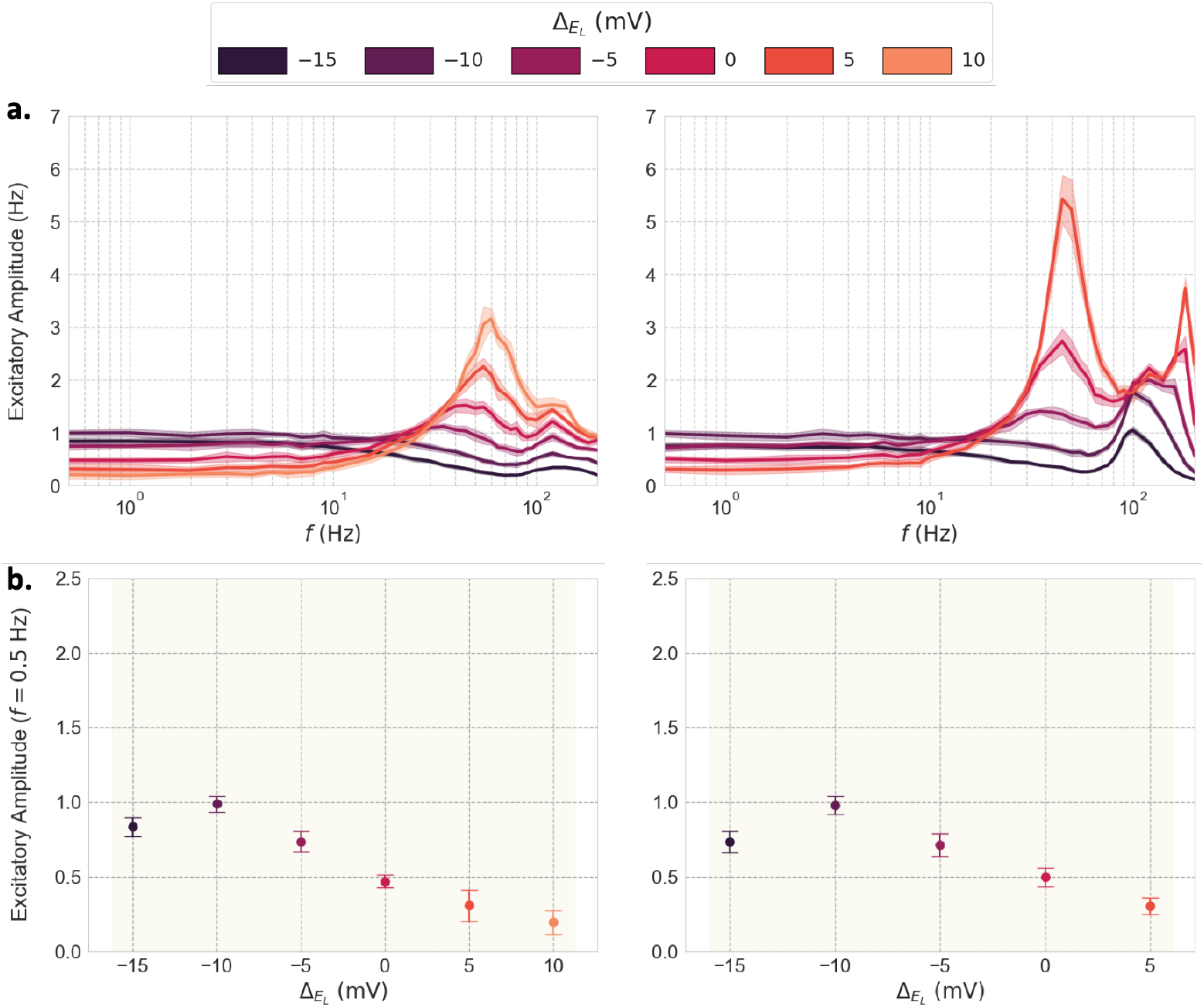
Evoked excitatory responses across 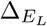 values (*ν*_drive_ = 4 Hz, *b* = 0 pA) for two choices of synaptic delay. **First column:** delay=1 ms; **second column:** delay=2 ms. **(a)** Evoked excitatory response amplitude versus input frequency. **(b)** Evoked excitatory response amplitudes to a slow input (*f* = 0.5 Hz).

### SI.9 Responsiveness results for changing 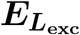

The equivalent results to those in Fig. 2 for 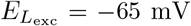 and 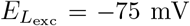 (as opposed to −70 mV) are shown in Fig. S9 and Fig. S10, respectively. Note the highly convergent responsiveness profiles for a given 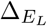 value (for a given *b*-*ν*_drive_ combination) across 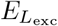 values.

**Fig. S9:**
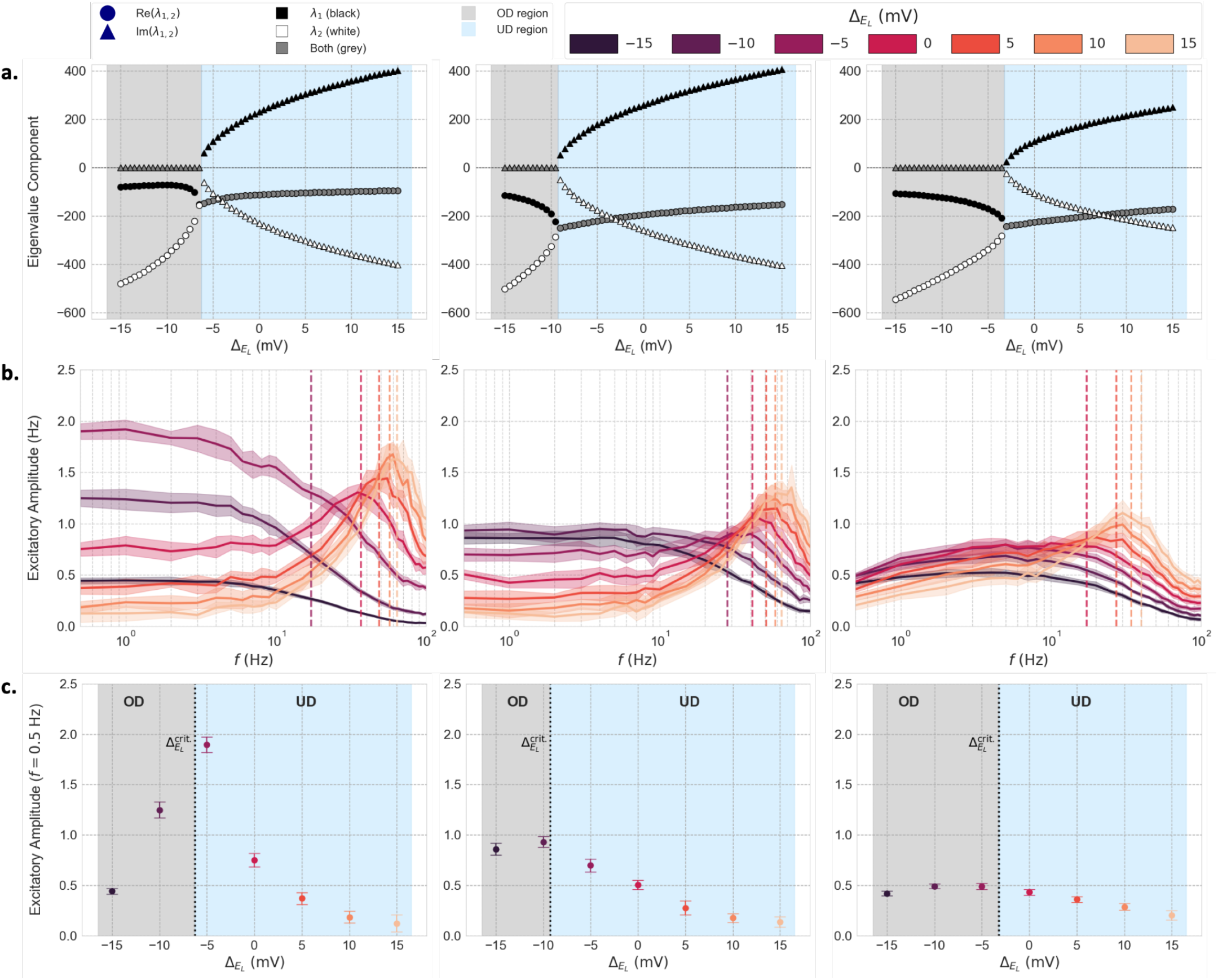
Mean-field linear stability analysis and SNN frequency-response profiles for different 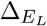 values 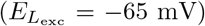 across parameter combinations (input amplitude *A* = 1 Hz). Columns and rows as in Fig. 2.

**Fig. S10:**
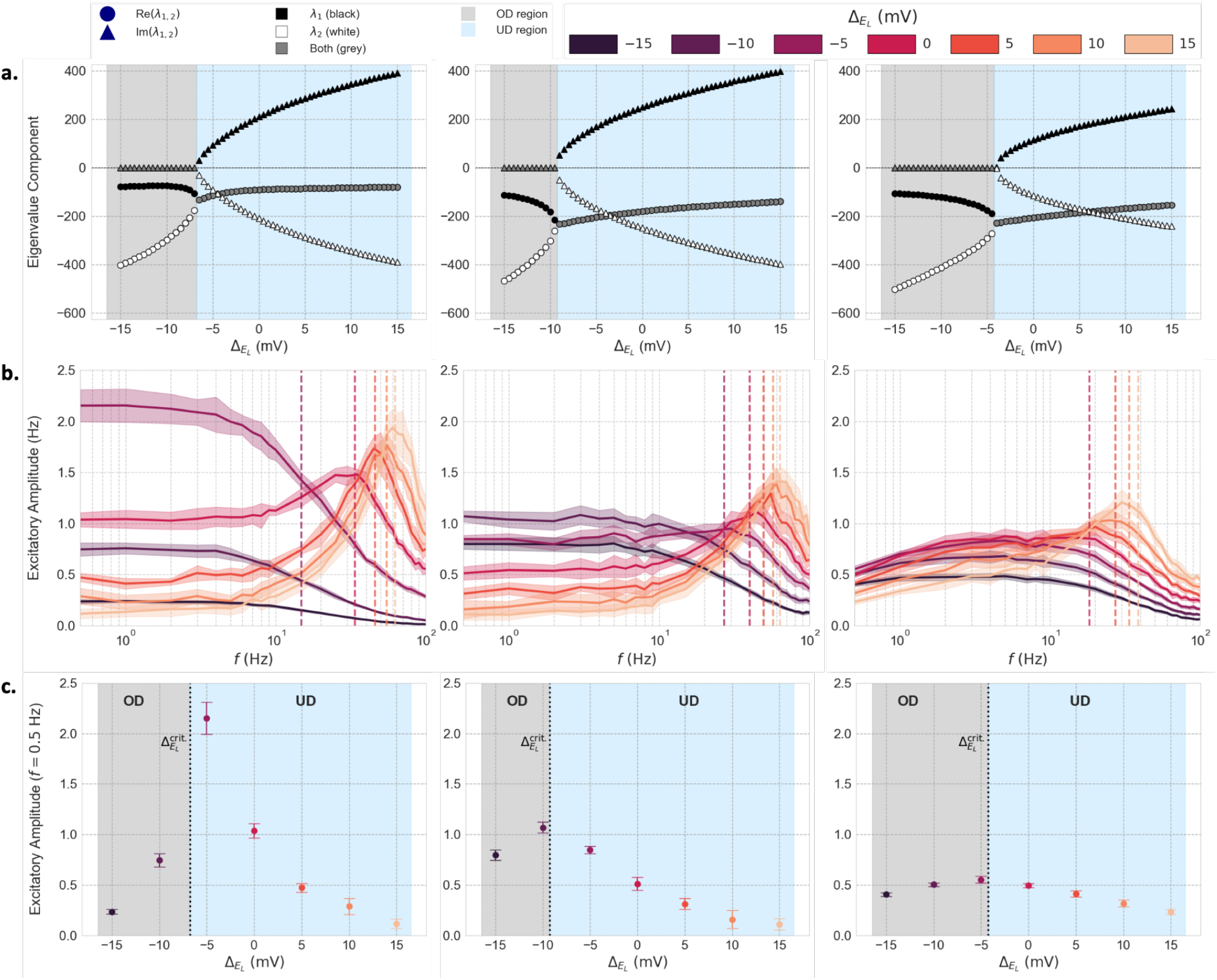
Mean-field linear stability analysis and SNN frequency-response profiles for different 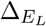 values 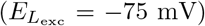 across parameter combinations (input amplitude *A* = 1 Hz). Columns and rows as in Fig. 2.

### SI.10 Fluctuation-normalised responses

When normalised against the standard deviations of the baseline fluctuations of their respective networks, the differential responsiveness properties of the OD and UD regimes are brought into sharper focus. As long as the evoked response is significantly larger than the baseline fluctuations, the stimulus is detected by the network, with the number of evoked spikes proportional to the amplitude of the response. However, if the evoked response is small relative to the fluctuations (≪ 1 std), the stimulus will not be registered.

For an input amplitude of *A* = 1 Hz (Fig. S11a), OD states yielded large evoked response amplitudes relative to their baseline fluctuations for low frequencies, up to the boundary of the resonant region, where their responsiveness steeply decreased. Conversely, UD states showed small evoked responses relative to baseline noise for all frequencies apart from a narrow frequency band surrounding their resonant frequencies. In this region, UD responses showed responsiveness greater than or equal to that of OD states, both in absolute and relative terms. Therefore, for sufficiently small inputs, UD networks function as precise frequency filters that are effectively unresponsive at all but their resonant frequency bands, at which they show stronger responses than all other networks in the parameter space.

For a larger input amplitude (*A* = 2 Hz, Fig. S11b), the same general effect was observed, albeit with UD networks showing detectable responses (∼ 1 std) across wider frequency bands. Unlike the *A* = 1 Hz case, where the responsiveness to slow inputs accorded with the analytical prediction (see 2.2), in the *A* = 2 Hz case, the input was sufficiently large to perturb networks far away from their operating points; for an amplitude of this size, the Taylor series approximation for the transfer function is less accurate. While most frequency-response profiles were qualitatively unchanged, a network close to the damping boundary showed both UD and OD characteristics (resonance peak and high low-frequency responsiveness, respectively; see Fig. S11b, column 1,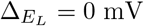.).

In summary, the fluctuation-adjusted responses emphasise the dual processing characteristics of OD and UD states, acting as broadband low-pass filters and narrowband high-frequency filters, respectively. In the latter case, the range of detectability correlates with the amplitude of the stimulus.

**Fig. S11:**
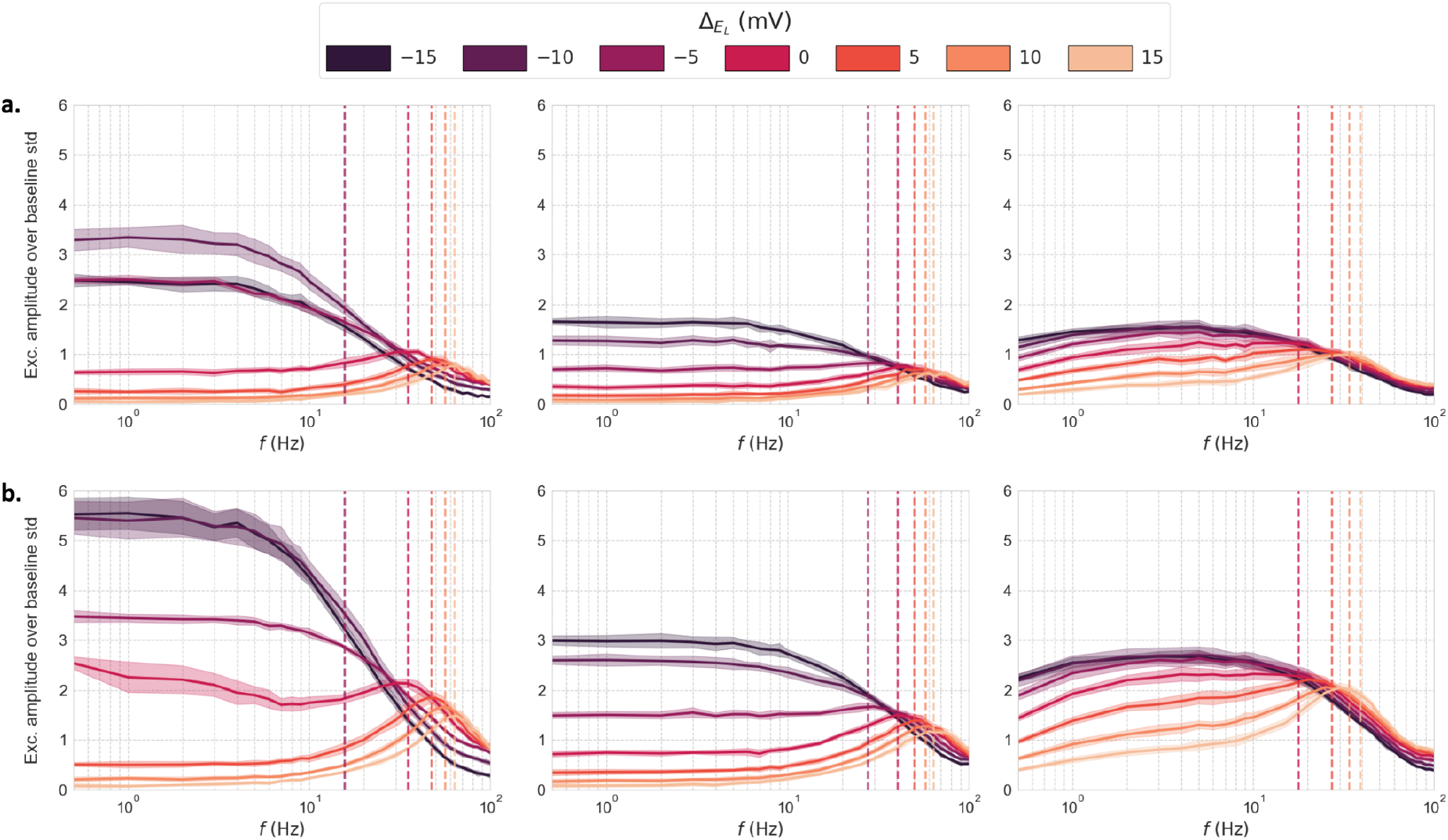
Evoked excitatory responses versus input frequency across 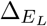 values, normalised by standard deviation of baseline excitatory firing rate fluctuations. **(a)** Input amplitude *A* = 1 Hz. **(b)** Input amplitude *A* = 2 Hz. **Columns 1-3**: parameter sets as in Fig. 2.

### SI.11 Transfer function derivation

#### SI.11.1 Subthreshold moments

A derivation of the subthreshold moments *µ*_*V*_, *σ*_*V*_ and *τ*_*V*_, following Di Volo et al. [15], is given below. First, the means and standard deviations of the excitatory and inhibitory conductances are computed as a function of the instantaneous excitatory and inhibitory presynaptic input rates, *ν*_*e*_ and *ν*_*i*_:

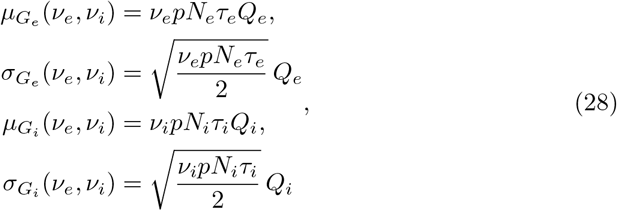

where *p* is the probability of connection between neurons, *N*_*e*_ (*N*_*i*_) is the size of the excitatory (inhibitory) population, *Q*_*e*_ (*Q*_*i*_) is the quantal conductance for excitatory (inhibitory) synapses and *τ*_*e*_ (*τ*_*i*_) is the excitatory (inhibitory) synaptic decay constant. The total synaptic conductance and the effective membrane time constant 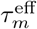 follow from these quantities:

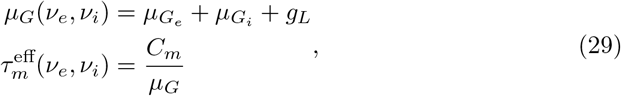

where *g*_*L*_ is the leak conductance and *C*_*m*_ is the membrane capacitance. From these quantities, the following expression for *µ*_*V*_, *σ*_*V*_ and *τ*_*V*_ are obtained (with the latter two taken directly from Zerlaut et al. [32]):

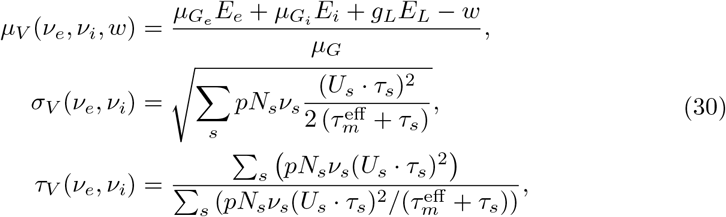

where *w* is the adaptation current, *E*_*e*_ (*E*_*i*_) is the excitatory (inhibitory) synaptic reversal potential, 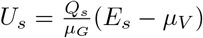 for *s* ∈ {*e, i*}.

#### SI.11.2 Fitting of effective threshold *V* ^eff^

The effective threshold is estimated as a second-order polynomial function of the subthreshold moments as follows [17]:

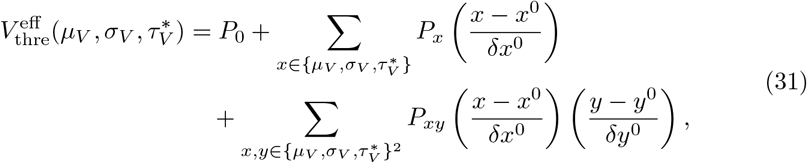

where 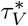 is a dimensionless expression for the autocorrelation time constant 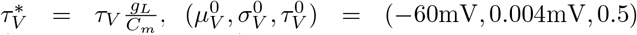 and 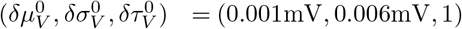 . A numerical fitting of the coefficients {*P*} was performed for each of the two cell types (RS and FS), yielding the values shown in Table S1.

**Table S1:** Fit parameters expressed in mV (from Di Volo et al. [15])

| Cell type | $P_0$ | $P_{\mu_V}$ | $P_{\sigma_V}$ | $P_{\tau_V^N}$ | $P_{\mu_V^2}$ | $P_{\sigma_V^2}$ | $P_{(\tau_V^N)^2}$ | $P_{\mu_V \sigma_V}$ | $P_{\mu_V \tau_V^N}$ | $P_{\sigma_V \tau_V^N}$ |
| --- | --- | --- | --- | --- | --- | --- | --- | --- | --- | --- |
| **RS** | -49.8 | 5.06 | -25 | 1.4 | -0.41 | 10.5 | -36 | 7.4 | 1.2 | -40.7 |
| **FS** | -51.4 | 4.0 | -8.3 | 0.2 | -0.5 | 1.4 | -14.6 | 4.5 | 2.8 | -15.3 |

